# Detecting Antibody Reactivities in Phage ImmunoPrecipitation Sequencing Data

**DOI:** 10.1101/2022.01.19.476926

**Authors:** Athena Chen, Kai Kammers, H Benjamin Larman, Robert B. Scharpf, Ingo Ruczinski

## Abstract

Phage ImmunoPrecipitation Sequencing (PhIP-Seq) is a recently developed technology to assess antibody reactivity, quantifying antibody binding towards hundreds of thousands of candidate epitopes. The output from PhIP-Seq experiments are read count matrices, similar to RNA-Seq data; however some important differences do exist. In this manuscript we investigated whether the publicly available method edgeR^1^ for normalization and analysis of RNA-Seq data is also suitable for PhIP-Seq data. We find that edgeR is remarkably effective, but improvements can be made and introduce a Bayesian framework specifically tailored for data from PhIP-Seq experiments (Bayesian Enrichment Estimation in R, BEER).

## Introduction

Because of their high abundance, easy accessibility in peripheral blood, and relative stability ex vivo, antibodies serve as excellent records of environmental exposures and immune responses. While several multiplexed methods have been developed to assess antibody binding specificities, Phage Immuno-Precipitation Sequencing (PhIP-Seq) is the most efficient technique available for assessing antibody binding to hundreds of thousands of peptides at cohort scale^2–4^. PhIP-Seq uses oligonucleotide library synthesis to encode proteome spanning peptide libraries for display on bacteriophages. These libraries are immunocaptured using an individual’s serum antibodies, and the antibody-bound library members are identified by high throughput DNA sequencing. The VirScan^4^ assay uses the PhIP-Seq method to quantify antibody binding to around 100,000 peptides spanning the genomes of more than 200 viruses that infect humans. Other commonly used libraries include the AllerScan^5^ and ToxScan libraries^6^, and a focused library for coronaviruses, including SARS-CoV-2^7^.

We and others have utilized PhIP-Seq to successfully identify novel autoantigens associated with autoimmune diseases^8–10^, to broadly characterize allergy-related antibodies^5^, to quantitatively compare the antibody repertoires of term and preterm neonates^11^, to assess changes in the anti-viral antibody response after bone marrow transplant^12^, to characterize the self-reactivity of broadly neutralizing HIV antibodies^13–15^, to link enteroviral infection with acute flaccid myelitis^16^, and for use in large crosssectional and longitudinal studies of exposure and response to hundreds of human viruses and thousands of bacterial proteins in healthy individuals and in individuals infected with HIV or measles^4,17–19^. In addition, we recently used PhIP-Seq to assess how antibody responses to endemic coronaviruses modulate COVID-19 convalescent plasma functionality^7^ and evaluated the heritability of antibody responses^20^.

The output from PhIP-Seq experiments are read count matrices, similar to RNA-Seq data, but important differences in the data structures, experimental design, and study objectives exist between the two sequencing-based methods. RNA-Seq experiments typically focus on differentially expressed genes or transcripts between experimental groups, rather than identifying expressed genes for any particular sample. The objective of PhIP-Seq experiments however is typically just that: detecting peptide antibody reactivity in an individual sample. Thus, in contrast to RNA-Seq experiments, the design of PhIP-Seq experiments requires the use of negative controls (i.e. “mock” immunoprecipitations (IPs) lacking antibody input, also referred to as beads-only samples), which are typically included as 4 to 8 wells of a 96-well plate. This generates a “n versus 1” mock IPs versus sample comparison, in contrast to the most common n_1_ versus n_2_ two-group comparison in RNA-Seq. In addition, genes with low read counts are presumed to have little biological relevance, and RNA-Seq data workflows typically filter out lowly expressed genes (measured as counts-per-million) prior to analysis. In PhIP-Seq experiments however, peptides with low read counts may have biological relevance and are not filtered out in advance. That said, under suitable assumptions, such as equality of variances in both groups, a two-group comparison with a single sample in one group can still be carried out.

Significant advances in normalization and analysis methods for RNA-Seq data have been made in recent years, with edgeR, DESeq2, and voom among the most popular open-source software packages available^1,21–23^. These methods model the number of reads using a negative binomial distribution to account for the inflated variance due to biological variability between samples in comparison to the expected variance of the binomial distribution. Parameter estimation is based on empirical Bayes methods to borrow strength across transcripts, stabilizing the estimates of the respective standard errors. Upregulated genes in RNA-Seq experiments draw a higher proportion of reads than expected for a given library size (here, total read counts), resulting in lower than expected read counts for other genes given that library size (‘‘competing resources”). Thus, a normalization factor for each sample is calculated in RNA-Seq experiments to account for this effect^24^. One assumption is that the majority of genes are not differentially expressed when comparing cases to controls.

In this manuscript we investigated whether the publicly available method edgeR^1^ for normalization and analysis of RNA-Seq data is also suitable for PhIP-Seq data. We highlight some of the differences between PhIP-Seq and RNA-Seq experiments and data sets, which motivates the development of a new methodology for PhIP-Seq data explicitly based on the assumed data generating mechanism, rather than adapting existing RNA-Seq approaches. To that end, we introduce a Bayesian framework specifically tailored for data from PhIP-Seq experiments (Bayesian Enrichment Estimation in R, BEER). Using simulation studies and data sets from existing HIV and SARS-CoV-2 studies, we investigate what improvements in sensitivity and specificity can be made, highlight the importance of empirical Bayes methods, and assess the effect of the number of mock IP samples on sensitivity and specificity.

## Results

### Simulation

Regardless of the approach used to estimate prior parameters, BEER has high discriminatory power for identifying enriched peptides (Figure 1, Supplementary Figure S1). In general, BEER using methods of moments (MOM) or maximum likelihood estimates (MLE) for the shape parameters in the beads-only prior distributions performed worse than BEER using edgeR parameter estimates, highlighting the importance of borrowing strength across peptides for improved parameter estimation (Supplementary Figure S1, Supplementary Table S1). The stability of parameter estimates also affected the improvement in BEER predictive performance by the number of beads-only samples used. While BEER with MOM and MLE parameter estimates greatly benefited from the inclusion of more beads-only samples in the experiment, BEER using edgeR parameter estimates had much less pronounced improvements as the number of beads-only samples was increased (Supplementary Figures S2, S3, and Supplementary Table S1). Using these edgeR parameter estimates, BEER posterior probabilities of enrichment were well-calibrated (Supplementary Figure S4), and estimates of fold changes were accurate (Supplementary Figure S5). Thus, we recommend the edgeR parameter estimates as default and imply their use when simply referring to BEER as the method used.

**Figure 1:**
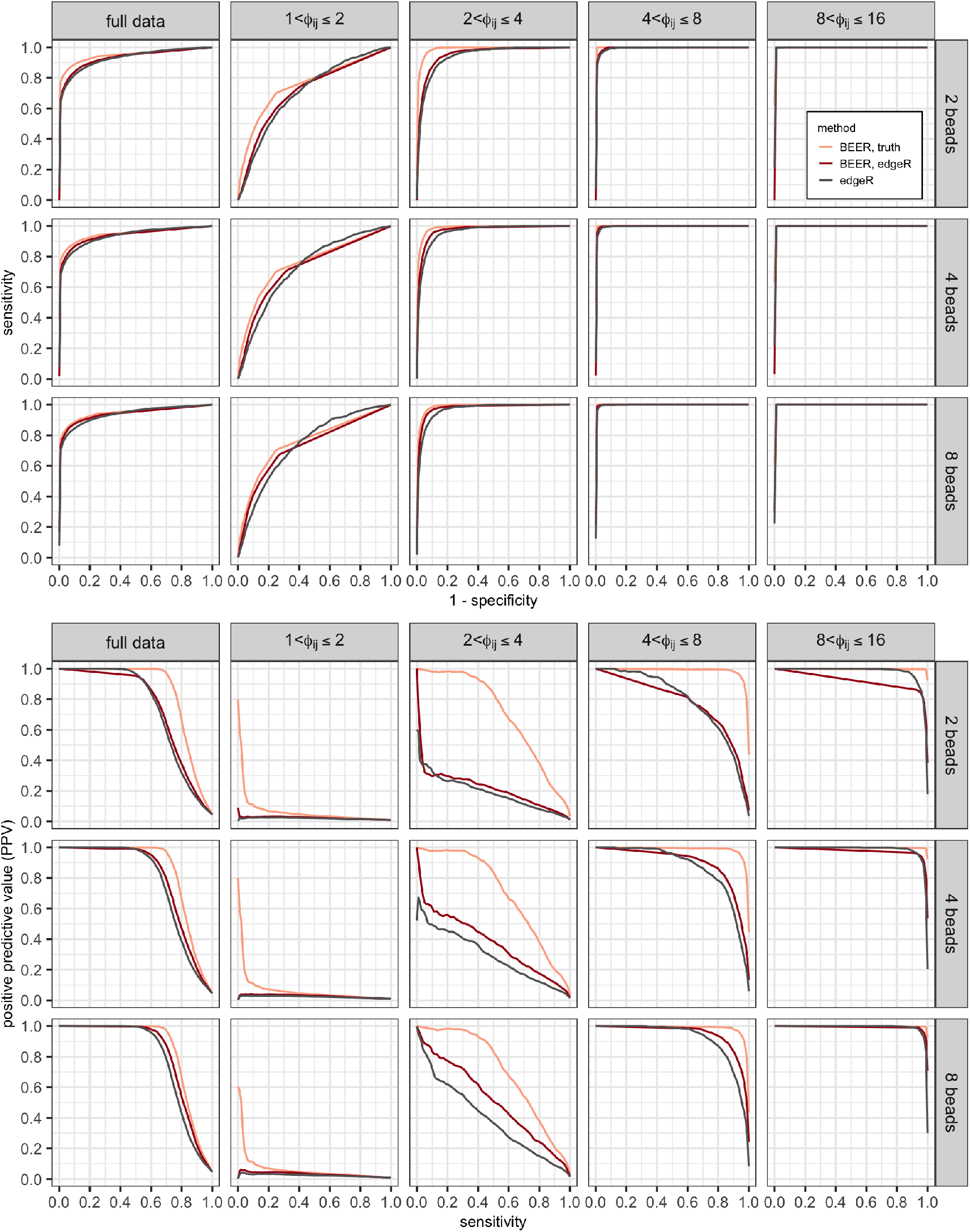
Average receiver operating characteristic (ROC; top panels) and precision-recall (PR; bottom panels) curves calculated from ten simulations, comparing edgeR (black lines) and BEER (red lines) across fold-change categories and number of beads-only samples available. Curves for BEER using the actual simulation shape parameters in the prior distributions (orange lines) are added to show the effect of sampling variability in these parameters. Results for fold changes above 16 are omitted since in all instances peptides were correctly classified as enriched.

In general, performances of edgeR and BEER for identifying enriched peptides were surprisingly similar (Figure 1, Supplementary Table S1). Both methods yielded near perfect receiver operating characteristic (ROC) curves for peptide fold changes above 4 (area under the curves (AUCs) > 0.99), and still outstanding ROC curves for fold changes between 2 and 4 (AUCs between 0.94 and 0.98) even when a design with only 2 beads-only samples was employed. Peptide fold changes less than 2 were harder to detect, reflected in substantially lower AUCs between 0.71 and 0.74. Precision-recall (PR) curves for edgeR and BEER are noticeably different for intermediate fold changes between 2 and 4, where also most improvement is (theoretically) possible for BEER if improved estimates of shape parameters used in the prior distributions were available. As expected from the near perfect ROC curves, reliable detection of peptides with fold changes above 4 is possible with a low rate of false positives. (Figure 1, Supplementary Table S2). For small fold changes less than 2 the positive predictive value (PPV) is generally poor, which is expected as the ROCs are modest and most peptides are not enriched.

Under commonly employed false discovery rate (FDR) control, BEER has a higher probability of correctly identifying enriched peptides than edgeR across all fold-changes, and the difference in probability is most pronounced for moderate fold changes between 2 and 8 (Figure 2). For example, under a FDR control of 5%, on average, the probability of identifying a peptide with a 4 fold change is 53% for BEER, but only 21% for edgeR. Similarly, BEER has a probability of at least 50% to detect fold changes above 3.7 under this FDR control, while edgeR requires a fold-change of at least 5.5. Of note, the BEER posterior probability cut-offs in the ten simulations to achieve a 5% FDR (see Methods) were between 0.25 and 0.49; thus, *using a commonly employed posterior probability cut-off of 0.5 leads to fewer false positives on average*.

**Figure 2:**
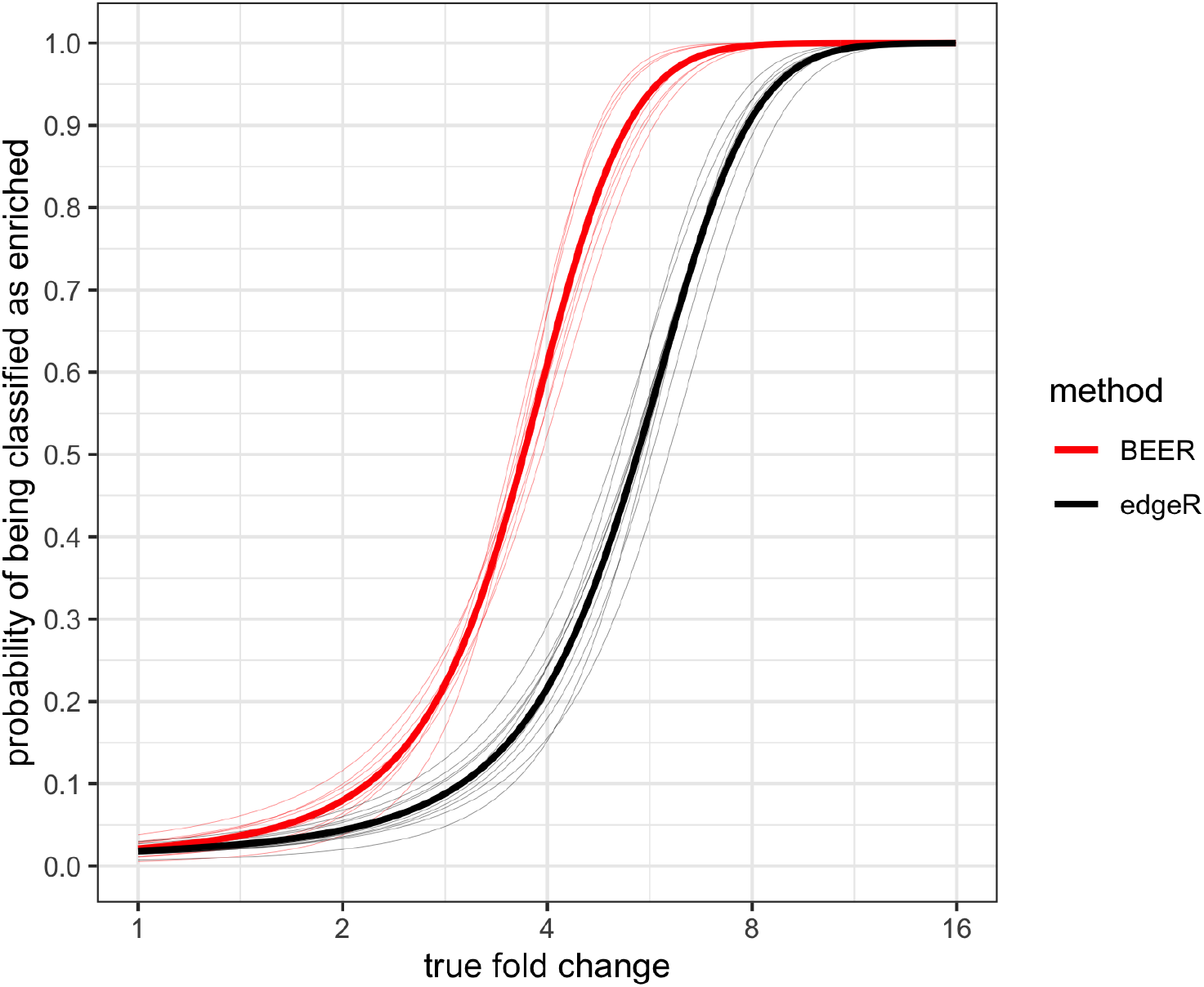
Estimated probabilities for correctly identifying enriched peptides (y-axis) as a function of the fold-change (x-axis) for each of ten simulated data sets based on logistic regression models. BEER posterior probability cut-offs were selected to achieve a false discovery rate of 5% in each simulation (see Methods). Thin lines indicate the individual simulations, thick lines the respective averages.

### HIV elite controllers

Both BEER and edgeR had no false positives across the six mock IP samples using a posterior probability cut-off of 0.5 for BEER and an FDR control of 5% for edgeR (corresponding to p-value cut-offs between 1.0 × 10^−3^ and 2.4 × 10^−3^ across the eight samples). As the non-replicated serum samples are from individuals infected with HIV subtype B, we expected stronger antibody reactivity to proteins from HIV subtype B, and indeed, BEER and edgeR detect more enrichments to peptides tiling proteins from HIV subtype B than proteins of any other HIV strain represented in the library (Figure 3). Notably, for any particular subtype B protein, BEER detects more enriched peptides than edgeR (while expected to have a lower type I error with a posterior probability cut-off of 0.5, see above). Some antibody reactivity to proteins from other HIV subtypes is expected due to cross-reactivity (Supplementary Figure S6).

**Figure 3:**
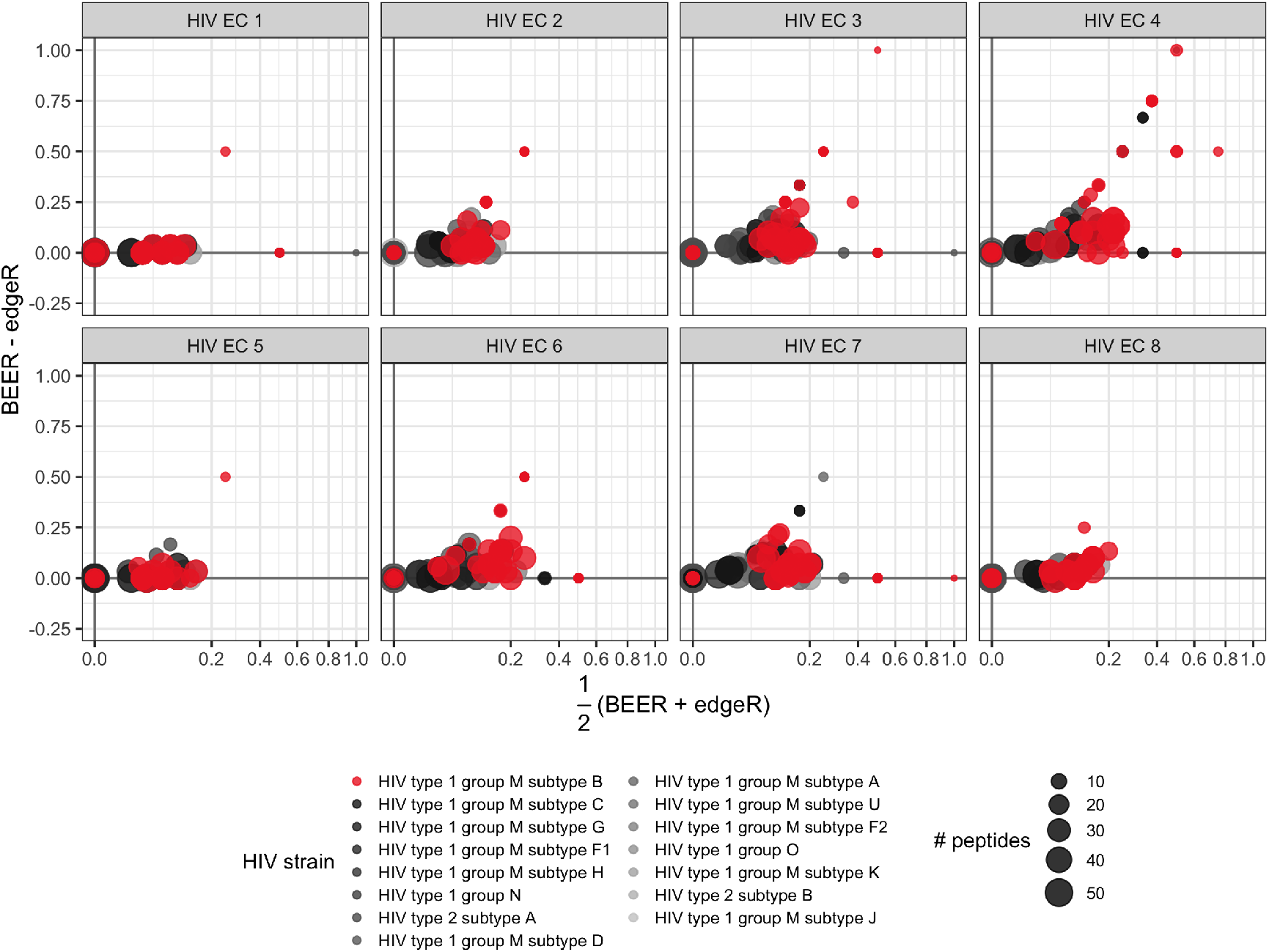
Bland-Altman (MA) plots for the proportion of enriched peptides by protein, for eight elite controller samples. Points represent individual proteins, point colors indicate protein virus types, point diameters indicate the number of peptides tiling the respective proteins. All subjects shown here were infected with subtype B (red).

BEER and edgeR detected enrichments generally agreed across two technical replicates of a sample from an HIV subtype A infected individual (Supplementary Figure S7). Using the same cut-offs as above for declaring enrichment, both methods had high agreement (concordance about 0.90–0.95) between the two samples among the top 100 peptides ranked by posterior probabilities and p-values, respectively (Supplementary Figure S8). This is not surprising as the BEER posterior probabilities for peptides ranked 100 in the two technical replicates were 0.99 and 1.00 respectively, and the edgeR p-values were 3.1 × 10^−4^ and 9.4 × 10^−5^ with respective 5% FDR cut-offs of 1.5 × 10^−3^ and 1.8 × 10^−3^, respectively. Thus, both methods exhibit high confidence that most of the peptides among the top 100 are truly enriched (Supplementary Figure S9). While BEER concordance decreases but remains above 0.90 when considering the lists of peptides with ranks up to 200, the edgeR concordance does drop more noticeably (Supplementary Figure S8), potentially indicating a higher sensitivity in BEER for peptides with smaller fold changes. Comparing subtype A peptides across technical replicates, BEER and edgeR had very similar performance. Compared to other subtypes, both methods also showed less discordance among subtype A peptides (Supplementary Table S3).

### CoronaScan

In a round-robin, leaving one mock IP sample out in turn, no false positives (i.e., peptides falsely called enriched) were produced by BEER or edgeR across the eight mock IPs in the CoronaScan data using a posterior probability cutoff of 0.5 for BEER and an FDR control of 5% for edgeR (p-value cutoffs ranged from 3.3 × 10^−4^ to 1.1 × 10^−3^). Among the six serum samples from individuals prior to the COVID-19 pandemic, BEER and edgeR show more enrichment to peptides tiling human coronaviruses, but generally no enrichment of SARS-CoV-2 peptides (VRC 1 – VRC 6, Supplementary Figure S10). In contrast, among the four samples from a single individual infected with SARS-CoV-2, taken at 10, 11, 12 and 13 days since symptom onset, an enrichment of SARS-CoV-2 protein tiling peptides is apparent. Particularly on day 13 after symptom onset, the patient presumably has produced a large number of antibodies which were detected by both BEER and edgeR. Of note, this was the first day the a SARS-CoV-2 antibody test was positive (D13, Supplementary Figure S10), further demonstrating the power and utility of the PhIP-Seq approach.

Comparing peptide replicates in the CoronaScan library, concordance among the most enriched peptides (when ranked by posterior probabilities and p-values for BEER and edgeR, respectively) was generally above 0.80 (Supplementary Figure S11). For example, in sample VRC 4 both methods show high confidence that the top 35 peptides are truly enriched (posterior probabilities of 0.97 and 1.00 for BEER, and p-values of 8.6 × 10^−5^ and 1.9 × 10^−6^ for edgeR, well below the 5% FDR cut-off derived p-value of 4.2 × 10^−4^, Supplementary Figure S12). BEER and edgeR perform very similarly in this example (with BEER slightly better between peptides 10 and 20), showing concordance above 0.80 before dropping significantly after about 50 peptides. The somewhat lower concordances compared to the same ranking metric derived from the two technical replicates in the HIV elite controllers above can possibly explained by the smaller range of proportions of reads pulled in the CoronaScan platform, and therefore higher correlation among these proportions comparing the technical replicates in the HIV data (Supplementary Figure S8 left, versus Supplementary Figure S11 left). Among all CoronaScan samples the overall concordance of peptide pair enrichment calls was outstanding, with less than 1% discordant calls in each sample, for both BEER using a posterior probability cutoff of 0.5 and 5% FDR cutoffs for edgeR (corresponding to p-value cutoffs between 3.3 × 10^−4^ and 1.1 × 10^−3^, Supplementary Table S4).

## Discussion

In this manuscript we investigated whether the publicly available method edgeR^1^ for normalization and analysis of RNA-Seq data is also suitable for PhIP-Seq data. With the exception of calculating one-sided p-values to infer peptide reactivity, no “tweaks” were necessary in the implementation, and we found the approach to be effective. However, using simulation studies we showed that substantial improvements are possible with a Bayesian framework specifically tailored for data from PhIP-Seq experiments (Bayesian Enrichment Estimation in R, BEER). In particular for peptides showing weaker reactivity, we saw an improvement of sensitivity with lower false positive rates when standard cut-offs were employed (posterior probability > 0.5 for the Bayesian method and a Benjamini-Hochberg false discovery rate control of 5% for edgeR). This comparison might be perceived as somewhat unfair, as the data were simulated from a model similar to that underlying BEER, which we recognize. However, BEER was implemented in a way we believe reflects the true data generating mechanism, which is also corroborated by the fact that BEER showed better performance also on real data such as the data from the HIV elite controllers, where BEER detected more enriched peptides of the correct HIV subtype than edgeR. This improved performance comes at a price of increased computational cost. While edgeR delivers almost instantaneous results, the Markov chains underlying BEER are time consuming. However, since laboratory prep and sequencing are expensive and certainly take more time than running such Markov chains, we believe utilizing extra CPU time to run BEER, as shown in our workflow (https://github.com/athchen/beer_manuscript), may yield worthwhile additional discoveries.

It was initially surprising to us how well edgeR fared on PhIP-Seq data despite being designed specifically for RNA-Seq data. While important differences between PhIP-Seq and RNA-Seq data structures exist, as previously described, edgeR captures some of the most important effects that exist in both types of data. For example, unlike RNA-Seq, the PhIP-Seq experimental protocol requires the use of negative controls (i.e. samples with no serum) on a 96-well plate. The observed read counts mapped to the peptides among those negative controls show a very strong peptide-dependent bias in library representation and/or “background” binding to the beads, such that some peptides consistently draw a much higher proportion of reads than others (Supplementary Figure S13). However, in a “n versus 1” mock IPs versus sample comparison where inference is drawn for each peptide, these differences among peptides are similar to the biological variability observed between genes in RNA-Seq^25^. In addition, edgeR models read counts using a negative binomial distribution to account for larger than binomial variability between samples, an effect we also observe in PhIP-Seq data (Supplementary Figure S14). And while we expect reactive peptides in a serum sample to pull a large number of reads, and thus – after adjusting for library size – expect non-reactive peptides in a serum-sample to have fewer reads on average than the corresponding beads-only sample peptides (Supplementary Figure S15), the resulting attenuation constant in essence is the same as the scale factors derived from the trimmed mean of M-values approach in edgeR^1,24^.

Our findings also highlight the importance of empirical Bayes methods for parameter estimation. Methods of moments and maximum likelihood estimates for individual peptide prior distribution shape parameters performed substantially worse than those obtained by borrowing strength across peptides. By also using the true shape parameters in our simulations to assess sensitivity and specificity, we were able to demonstrate that, particularly for intermediate fold changes, better performance could be achieved by improving procedures to estimate these parameters. Our findings also give guidance for experimental design, such as the chosen number of mock IPs per 96-well plate. Allocating more beads-only samples to a plate improves estimation of these shape parameters that largely quantify between sample variability of the probabilities of a specific peptide to pull a read. Choosing more beads-only samples means reduced number of biological samples assayed per plate, which for the practitioner means additional cost and labor for more plates. In previous experiments, the number of mock IPs per plate was typically between 4 and 8. Our simulation studies showed that this is appropriate, as the observed difference in performance between 8 and 4 beads-only samples was much less than the observed difference between 4 and 2 mock IPs, indicating diminishing returns.

A few technical details should also be discussed further. As described in the Methods section, reads from highly reactive peptides (initial fold change estimate above 15) are removed from the data of mock IPs and the actual sample before BEER analysis, and the respective library sizes are recalculated. The main reason for doing so is simply to stabilize the inference and improve scalabilty, as allowing for extreme fold changes in the Bayesian model for a few peptides can affect these features. We verified that there were “no false positives” in the sense that all posterior probabilities for these highly reactive peptides were 1 if not excluded from the analyses. Thus, the chosen “highly enriched” threshold of 15 is likely conservative. We also note that the Bayesian model can be extended to run Markov chains for multiple samples against the beads-only samples. However, the resulting increase in parameter space makes this a challenging endeavor, especially with regards to scalability. It could be argued that for the same reasons stated above an increase in CPU time should be acceptable if this leads to an improvement detecting reactive peptides. However, we did not observe an improvement in detecting antibody reactivities in simulation studies we performed (data now shown). No notable improvements were observed when the same peptide was simulated as enriched in all samples compared to the beads-only, and a deterioration was observed when reactivity was not common to all peptides. Since in real life experiments we seldom expect the exact same peptides to be reactive, we did not pursue this line of research further.

In summary, antibodies commonly serve as indicators of environmental exposures and immune responses, and Phage ImmunoPrecipitation Sequencing allows for quantification of antibody binding to hundreds of thousands of peptides, in individuals and large cohorts. We believe that this technology will play an even more prominent role in the future, addressing questions about exposures and health outcomes in populations, as well as individualized medicine. In this manuscript, we introduce a method and a software package for analyzing data from this technology, contrast it with an existing RNA-Seq software package that can be retooled for PhIP-Seq data, and share a workflow with practitioners to successfully carry out their own analyses of data resulting from PhIP-Seq experiments.

## Methods

### A Bayesian model for detecting antibody enrichment

A succinct summary of the model notation is provided in Supplementary Table S5. On a 96-well plate suppose we observe *Y_ij_* read counts for peptide *i* ∈ {1, 2,…, *P*} in sample *j* ∈ {1, 2,…, 96}. Let *n_j_* = ∑_*i*_*Y_ij_* denote the total read count (library size) for sample *j*. Without loss of generality, assume samples {1, 2,…, *N*} are mock IP (beads-only) samples. To infer reactivity, we compare one sample to all beads-only samples on the same plate. Our hierarchical model to infer peptide reactivity in a sample *j* ∈ {*N* + 1,…, 96} is described as follows.

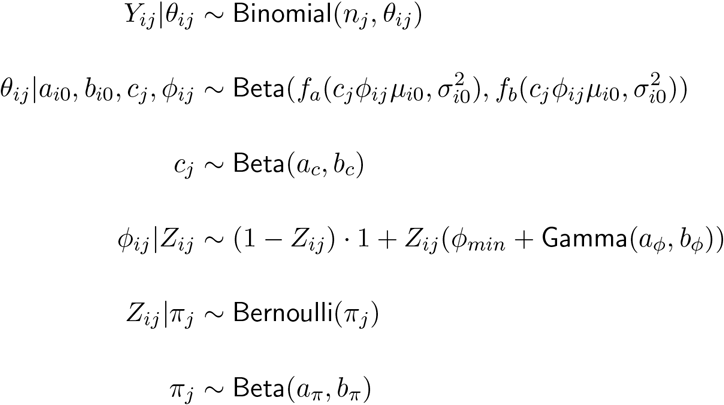

The main parameter of interest *Z_ij_* is a binary indicator denoting whether peptide *i* elicits an enriched antibody response in sample *j* (a 0 indicates no, a 1 indicates yes). The prior is a Bernoulli with success probability *π_j_*. For all mock IP samples, this success probability is zero. For sample *j*, *π_j_* is modeled as a beta distribution. The shape parameters *a_π_* and *b_π_* of the Beta prior distributions are chosen as 2 and 300 in our applications, to reflect peptide enrichment seen in previous studies, but also make it sufficiently diffuse to support a range of proportions (Supplementary Figure S16). The parameter *ϕ_ij_* is the fold change observed for peptide *i* in sample *j*. It is equal to 1 if *Z_ij_* = 0, i.e., peptide *i* does not elicit an enriched antibody response in sample *j*. Enriched peptides are expected to pull a larger proportion of reads, so only fold changes larger than 1 are considered. Here, we model the fold-change as a shifted gamma distribution (with shape parameters *a_ϕ_* = 1.25 and *b_ϕ_* = 0.1, Supplementary Figure S16), with the magnitude of the shift *ϕ_min_* being the minimum fold-change assumed for an enriched peptide (chosen as 1 in our applications). In the presence of reactive peptides pulling reads, non-reactive peptides in sample *j* will have less reads than expected from the beads-only samples where no reactive peptides exist by definition. We denote this attenuation constant for sample *j*, which is similar to the trimmed mean of M-values (TMM) scale factor used in edgeR^24^, with *c_j_*. Typically, only a minority of peptides in a sample show reactivity and the attenuation constant usually is between 0.5 and 1 (being equal to 1 in mock IP samples). In this application, we chose a Beta prior with scaling constants *a_c_* = 80 and *b_c_* = 20 (Supplementary Figure S16; the attenuation constant is equal to 1 in the mock IP samples). The observed read counts *Y_ij_* are modeled using a Binomial(*n_j_*, *θ_ij_*) distribution, where *θ_ij_* denotes the probability that peptide *i* pulls a read in sample *j*, and *n_j_* denotes the total library size in sample *j*. This Binomial probability is modeled using a Beta prior distribution, and the shape parameters depend on the expected peptide read counts observed in the mock IP samples (estimation procedures described below), the fold change *ϕ_ij_*, and the attenuation constant based on the reads pulled by all reactive peptides in the sample.

### Shape parameter estimation

We define two functions, *f_a_* and *f_b_*, used for the description of the Beta shape parameters *a* and *b* given a mean *μ* and a variance *σ*^2^.

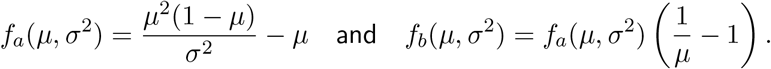

The mean and variance for peptide *i* in a beads-only sample (e.g., for a Beta distribution with shape parameters *a*_*i*0_ and *b*_*i*0_) is given by

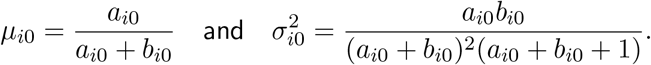

Since each sample generally contains more than a million of reads, estimates of the Binomial probabilities 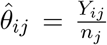 in the mock IP samples *j* ∈ {1, 2,…, *N*} are very precise. Method of moments (MOM) estimates for the peptide *i* shape parameters *a*_*i*0_ and *b*_*i*0_ can be derived by equating the mean and variance of the above Binomial estimates across all beads-only samples to the mean and variance of the Beta(*a*_*i*0_, *b*_*i*0_) distribution.

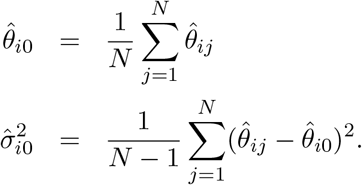

The MOM estimates for *a*_*i*0_ and *b*_*i*0_ are then given by

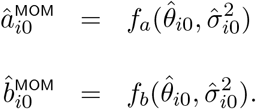

Maximum likelihood estimates (MLEs) for *a*_*i*0_ and *b*_*i*0_ were derived using the Broyden, Fletcher, Goldfarb and Shanno quasi-Newton optimiziation algorithm with box constraints^26^, as implemented in the R `optim()` function.

Numerous papers have demonstrated the benefits of shrinkage or variance stabilization in high throughput genomics experiments, borrowing strength across units such as genes and proteins^1,22,27,28^. This can be particularly important when the sample sizes are small, such as the number of mock IP experiments on each plate in our application, but neither the MLEs nor the MOM estimates described above make use of this. In contrast for example, edgeR uses an emprical Bayes approach^29^ to approximate the larger than binomial variability observed in the RNA-Seq read counts, and to stabilize these variance estimates, which are characterized by the tagwise dispersion parameter (the squared coefficient of variation of 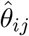, denoted here as 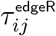)^1^. We note that we can use the estimates of these tagwise dispersion parameters to derive new estimates of the variances for our Binomial probabilities *θ_ij_*. Specifically, for peptide *i* we have 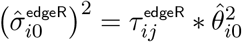, and thus

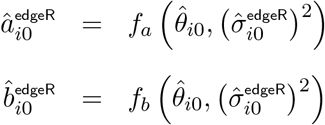

The Beta parameters *a* and *b* can be thought of as the number of successes and the number of failures, respectively, in *a* + *b* trials. The Markov Chain Monte Carlo (MCMC) sampler Just Another Gibbs Sampler (JAGS) can encounter numerical issues when either of those is less than 1. In PhIP-Seq experiments *a* is much smaller than *b* as a peptide only pulls a fraction of the total number of reads even when reactive. Thus, to avoid these numerical problems, we set *a* to be the larger number of the estimated value and 1, and then estimate *b*.

### Markov chain Monte Carlo

The model was implemented in JAGS (4.3.0) and run using the `R` interface for JAGS, `rjags`^30–32^. We use maximum likelihood estimates to select starting values of *θ_ij_*, *Z_ij_*, *ϕ_ij_*, *c_j_*, and *π_j_* to initialize the Markov chain for non beads-only sample *j*. As described above, 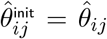 is the MLE of the binomial probability calculated from the read counts. Since *Z_ij_* is needed to update *c_j_*, *π_j_*, *ϕ_ij_*, we set 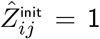 if its observed read count is at least twice as large as the expected read count in a beads-only sample. That is, 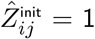 if 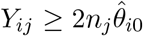, and 0 otherwise. The initial value for the attenuation constant is derived by regressing the observed read counts on the expected reads count for all non-enriched peptides in that sample, with 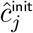 being the slope estimate

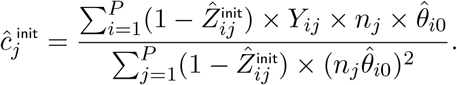

The initial value for the proportion of enriched peptides is the average of all enrichment indicators

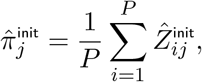

and the respective peptide fold changes are initialized as

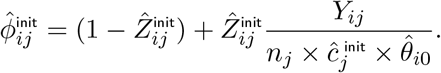

Since *c_j_* and *π_j_* are modeled using Beta distributions with no support at values 0 and 1, we use a small offset in the event that 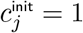 and 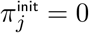.

In PhIP-Seq experiments we commonly observe very reactive peptides^18,33^. Allowing for extreme fold changes in the Bayesian model for a few peptides can affect the inference for other less reactive peptides, and can have negative consequences for numerical stability and scalability. In our applications, clearly enriched peptides defined as 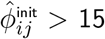 were filtered out before starting the Markov chain. Reads from such peptides in the mock IP and actual samples were removed, and the library sizes were recalculated.

### Peptide reactivity detection with edgeR

To identify reactive peptides, each serum sample is compared to all beads-only samples from the same plate. Differential expression in edgeR is assessed for each unit (here, each peptide) using an exact test analogous to Fisher’s comparing means between two groups of negative binomial random variables, but adapted for overdispersed data^34^. Two-sided p-values were subsequently converted to one-sided p-values as the alternative to the null of no reactivity (fold change = 1) is reactivity, leading to read count enrichment and thus, fold-changes larger than 1. Multiple comparisons corrections were based on the Benjamini–Hochberg procedure, using false discovery rates to delineate enrichment across all peptides.

### Simulation study

We simulated ten data sets based on the read counts observed in the HIV elite controller data described below. Each of these data sets had eight beads-only samples and twelve simulated serum samples. The twelve samples contain one beads-only sample run as an actual sample and two technical replicates (samples generated from the same parameters). For each simulated serum sample, we randomly selected 50 peptides as reactive. Among those, 10 peptides each had fold changes between 1 and 2, between 2 and 4, between 4 and 8, between 8 and 16, and between 16 and 32. Each data set was analyzed using the first two, four, and all eight beads-only samples to assess the sensitivity of the results to the number of beads-only used for analysis. For each data set and number of beads-only sample combination, we ran BEER with the true beads-only Beta *a*_0_, *b*_0_ prior parameters, estimated beads-only parameters using maximum likelihood, method of moments, and edgeR derived estimates.

Performance was primarily assessed using ROC and PR curves on the full data and fold-change subsets of the data. For each fold-change bin, curves were generated using all non-enriched peptides and enriched peptides within the specified fold-change group from the simulated serum samples (no peptides from beads-only samples were included). To ensure that all curves had the same support points, we used linear interpolation to approximate the sensitivity or positive predictive value respectively at each support point for each simulation. ROC curves started at 0 sensitivity and 0 false-positive rate, while PRC curves started at 0 sensitivity and perfect positive-predictive value. The interpolated curves were averaged point-wise to generate an average curve for each condition. The area under each ROC curve was calculated using trapezoidal approximation from the interpolated data points. We also used logistic regression to model the probability of identifying and enriched peptide by fold-change in each data set. Multiple comparisons for edgeR p-values were addressed using the Benjamini-Hochberg procedure to ensure a 5% FDR. Cut-offs for the posterior probabilities were selected in each data set to achieve 5% false positive calls.

### Examples

Antibody reactivity counts for eight plates of data were generated using the PhIP-Seq assay and the VirScan library on serum samples from HIV elite controllers with HIV subtype A and B infections, and analyzed by Kammers et al.^15^ to assess antibody profiles in HIV controllers and persons with treatment-induced viral suppression. We used count data for the 3,395 phage-displayed peptides spanning the HIV proteome in the VirScan library for ten samples and six-beads-only samples from one plate of data. Two of the ten samples are identical, run in duplicate on the same plate. To quantify the false-positive rate of each algorithm, we also ran each beads-only sample against the remaining five-beads-only samples in a round-robin.

The CoronaScan data consists of counts for 6,932 peptides for 10 serum samples and 8 beads-only samples from one plate of data^7^. Among the ten samples, six were pre-pandemic samples and four samples were from one individual infected with SARS-CoV-2. Samples from this individual were collected on days 10 through 13 since symptom onset. By design, each peptide is present in duplicate in the CoronaScan library, enabling us to assess the concordance of the fold-change estimates and the enrichment status within samples. We again ran each beads-only sample against the remaining 7 beads-only samples to assess false positive rates.

The example in the Discussion to highlight the strong peptide-dependent background binding to the beads was from a previous study to evaluates HIV antibody responses and their evolution during the course of HIV infection^18^ and to generate a classifier for recent HIV infections^33^.

## Acknowledgments

Funding for this work was provided by grants from the United States National Institutes of General Medical Science (NIGMS R01 GM136724) and Allergy and Infectious Diseases (NIAID R01 AI095068), and the National Cancer Institute (NCI P50 CA062924, NCI P30 CA006973).

## Software and Data

The data and code for the figures, tables, and benchmarks are freely available at https://github.com/athchen/beer_manuscript to ensure the reproducibility of our results. BEER can be run using the `R` package `beer` which is available on Github: https://github.com/athchen/beer (submitted to Bioconductor).

## Supplementary Materials

**Table S1:** Area under the ROC curves shown in Figure 1. Both BEER using edgeR parameter estimates and edgeR had near perfect classification for peptides with fold-changes above 4, even when only four beads-only samples were used to estimate *a*_*i*0_ and *b*_*i*0_.

|  | 2 beads-only | 4 beads-only | 8 beads-only |
| --- | --- | --- | --- |
| **BEER, MOM** | 0.884 | 0.939 | 0.944 |
| $1 < \phi_{ij} \leq 2$ | 0.695 | 0.757 | 0.756 |
| $2 < \phi_{ij} \leq 4$ | 0.881 | 0.959 | 0.975 |
| $4 < \phi_{ij} \leq 8$ | 0.937 | 0.989 | 0.994 |
| $8 < \phi_{ij} \leq 16$ | 0.953 | 0.994 | 0.995 |
| **BEER, MLE** | 0.886 | 0.939 | 0.944 |
| $1 < \phi_{ij} \leq 2$ | 0.697 | 0.759 | 0.762 |
| $2 < \phi_{ij} \leq 4$ | 0.886 | 0.959 | 0.976 |
| $4 < \phi_{ij} \leq 8$ | 0.940 | 0.989 | 0.994 |
| $8 < \phi_{ij} \leq 16$ | 0.953 | 0.994 | 0.996 |
| **BEER, edgeR** | 0.929 | 0.936 | 0.939 |
| $1 < \phi_{ij} \leq 2$ | 0.712 | 0.727 | 0.728 |
| $2 < \phi_{ij} \leq 4$ | 0.951 | 0.970 | 0.977 |
| $4 < \phi_{ij} \leq 8$ | 0.992 | 0.994 | 0.996 |
| $8 < \phi_{ij} \leq 16$ | 0.995 | 0.995 | 0.996 |
| **edgeR** | 0.926 | 0.934 | 0.936 |
| $1 < \phi_{ij} \leq 2$ | 0.709 | 0.728 | 0.737 |
| $2 < \phi_{ij} \leq 4$ | 0.940 | 0.960 | 0.967 |
| $4 < \phi_{ij} \leq 8$ | 0.991 | 0.994 | 0.995 |
| $8 < \phi_{ij} \leq 16$ | 0.996 | 0.996 | 0.996 |

**Table S2:**
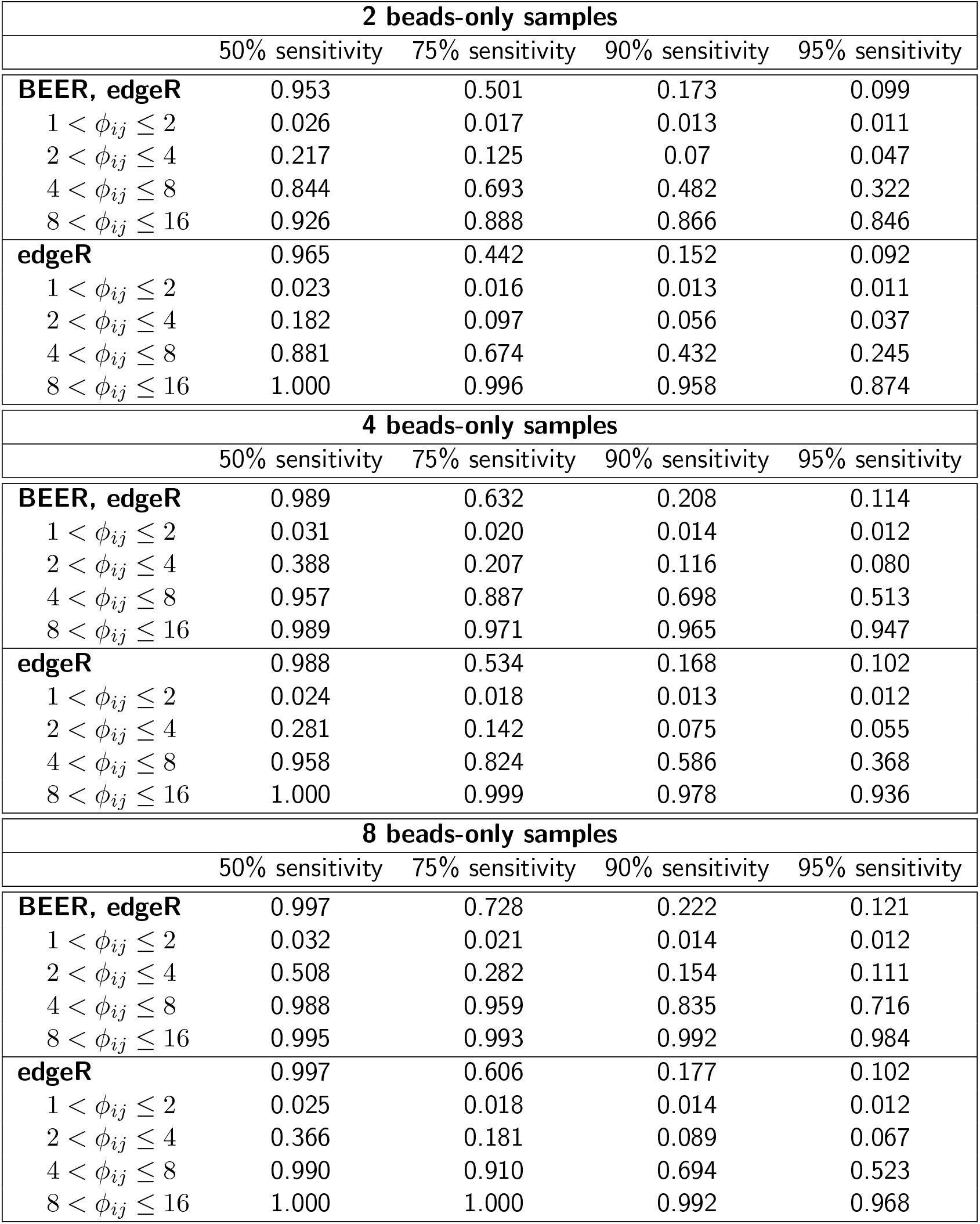
Average positive predictive values for select sensitivities for the curves in Figure 1.

| 2 beads-only samples |  |  |  |  |
| --- | --- | --- | --- | --- |
|  | 50% sensitivity | 75% sensitivity | 90% sensitivity | 95% sensitivity |
| **BEER, edgeR** | 0.953 | 0.501 | 0.173 | 0.099 |
| $1 < \phi_{ij} \leq 2$ | 0.026 | 0.017 | 0.013 | 0.011 |
| $2 < \phi_{ij} \leq 4$ | 0.217 | 0.125 | 0.07 | 0.047 |
| $4 < \phi_{ij} \leq 8$ | 0.844 | 0.693 | 0.482 | 0.322 |
| $8 < \phi_{ij} \leq 16$ | 0.926 | 0.888 | 0.866 | 0.846 |
| **edgeR** | 0.965 | 0.442 | 0.152 | 0.092 |
| $1 < \phi_{ij} \leq 2$ | 0.023 | 0.016 | 0.013 | 0.011 |
| $2 < \phi_{ij} \leq 4$ | 0.182 | 0.097 | 0.056 | 0.037 |
| $4 < \phi_{ij} \leq 8$ | 0.881 | 0.674 | 0.432 | 0.245 |
| $8 < \phi_{ij} \leq 16$ | 1.000 | 0.996 | 0.958 | 0.874 |
| 4 beads-only samples |  |  |  |  |
|  | 50% sensitivity | 75% sensitivity | 90% sensitivity | 95% sensitivity |
| **BEER, edgeR** | 0.989 | 0.632 | 0.208 | 0.114 |
| $1 < \phi_{ij} \leq 2$ | 0.031 | 0.020 | 0.014 | 0.012 |
| $2 < \phi_{ij} \leq 4$ | 0.388 | 0.207 | 0.116 | 0.080 |
| $4 < \phi_{ij} \leq 8$ | 0.957 | 0.887 | 0.698 | 0.513 |
| $8 < \phi_{ij} \leq 16$ | 0.989 | 0.971 | 0.965 | 0.947 |
| **edgeR** | 0.988 | 0.534 | 0.168 | 0.102 |
| $1 < \phi_{ij} \leq 2$ | 0.024 | 0.018 | 0.013 | 0.012 |
| $2 < \phi_{ij} \leq 4$ | 0.281 | 0.142 | 0.075 | 0.055 |
| $4 < \phi_{ij} \leq 8$ | 0.958 | 0.824 | 0.586 | 0.368 |
| $8 < \phi_{ij} \leq 16$ | 1.000 | 0.999 | 0.978 | 0.936 |
| 8 beads-only samples |  |  |  |  |
|  | 50% sensitivity | 75% sensitivity | 90% sensitivity | 95% sensitivity |
| **BEER, edgeR** | 0.997 | 0.728 | 0.222 | 0.121 |
| $1 < \phi_{ij} \leq 2$ | 0.032 | 0.021 | 0.014 | 0.012 |
| $2 < \phi_{ij} \leq 4$ | 0.508 | 0.282 | 0.154 | 0.111 |
| $4 < \phi_{ij} \leq 8$ | 0.988 | 0.959 | 0.835 | 0.716 |
| $8 < \phi_{ij} \leq 16$ | 0.995 | 0.993 | 0.992 | 0.984 |
| **edgeR** | 0.997 | 0.606 | 0.177 | 0.102 |
| $1 < \phi_{ij} \leq 2$ | 0.025 | 0.018 | 0.014 | 0.012 |
| $2 < \phi_{ij} \leq 4$ | 0.366 | 0.181 | 0.089 | 0.067 |
| $4 < \phi_{ij} \leq 8$ | 0.990 | 0.910 | 0.694 | 0.523 |
| $8 < \phi_{ij} \leq 16$ | 1.000 | 1.000 | 0.992 | 0.968 |

**Table S3:**
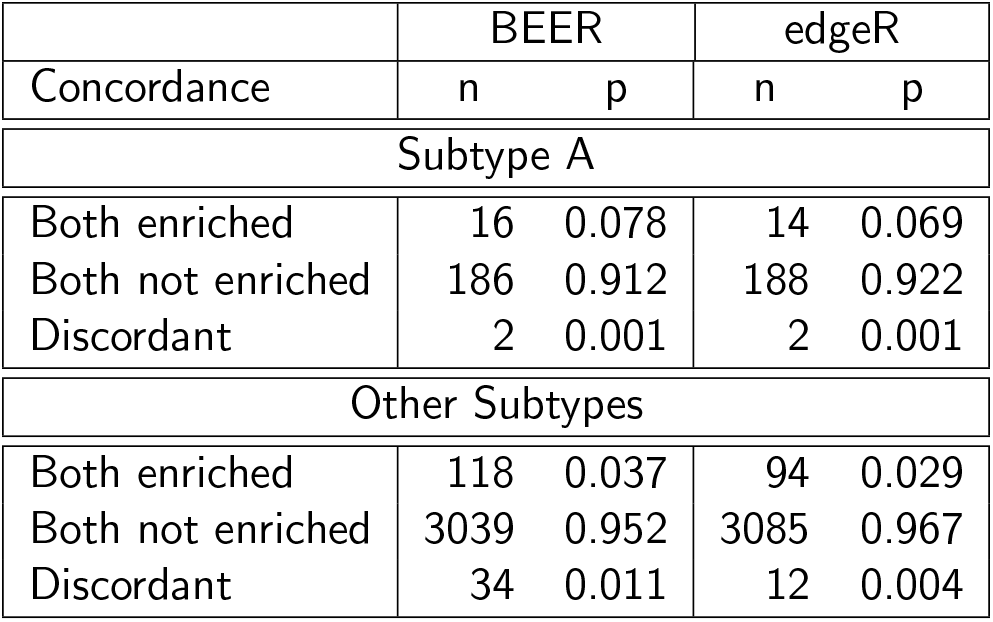
Concordance of enrichment calls between two technical replicates of an HIV subtype A infected individual for BEER and edgeR. A total of 204 peptides from subtype A and 3,191 peptides from other subtypes were present on the platform. n: number of peptides; p: proportion of peptides.

**Table S4:** Concordance of enrichment calls between peptide pairs for all CoronaScan samples. Each sample has 3,366 unique peptide pairs. n: number of pairs with concordant enrichment calls; p: proportion of pairs with concordant enrichment calls.

| Sample | BEER |  | edgeR |  |
| --- | --- | --- | --- | --- |
|  | n | p | n | p |
| VRC 1 | 3452 | 0.996 | 3454 | 0.997 |
| VRC 2 | 3443 | 0.993 | 3443 | 0.993 |
| VRC 3 | 3430 | 0.990 | 3423 | 0.988 |
| VRC 4 | 3432 | 0.990 | 3427 | 0.989 |
| VRC 5 | 3445 | 0.994 | 3444 | 0.994 |
| VRC 6 | 3435 | 0.991 | 3433 | 0.990 |
| SARS-CoV-2, D10 Ab- | 3455 | 0.997 | 3450 | 0.995 |
| SARS-CoV-2, D11 Ab- | 3450 | 0.995 | 3446 | 0.994 |
| SARS-CoV-2, D12 Ab- | 3441 | 0.993 | 3439 | 0.992 |
| SARS-CoV-2, D13 Ab+ | 3441 | 0.993 | 3437 | 0.992 |

**Table S5:**
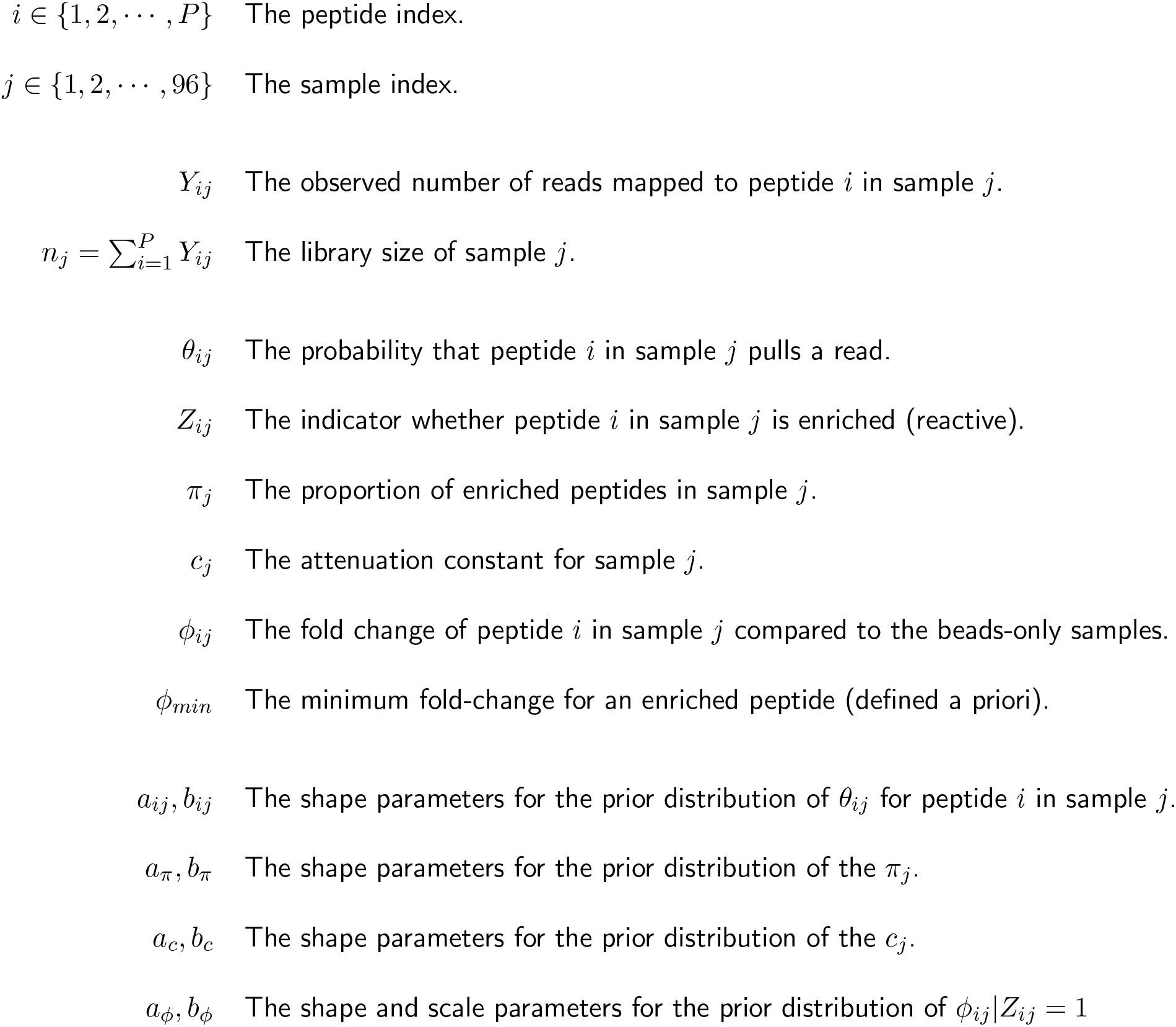
The notation used in the BEER model. Parameters specific to beads-only samples are denoted with the subscript *i*0 (e.g. *a*_*i*0_, *b*_*i*0_, *θ*_*i*0_, etc.).

**Figure S1:**
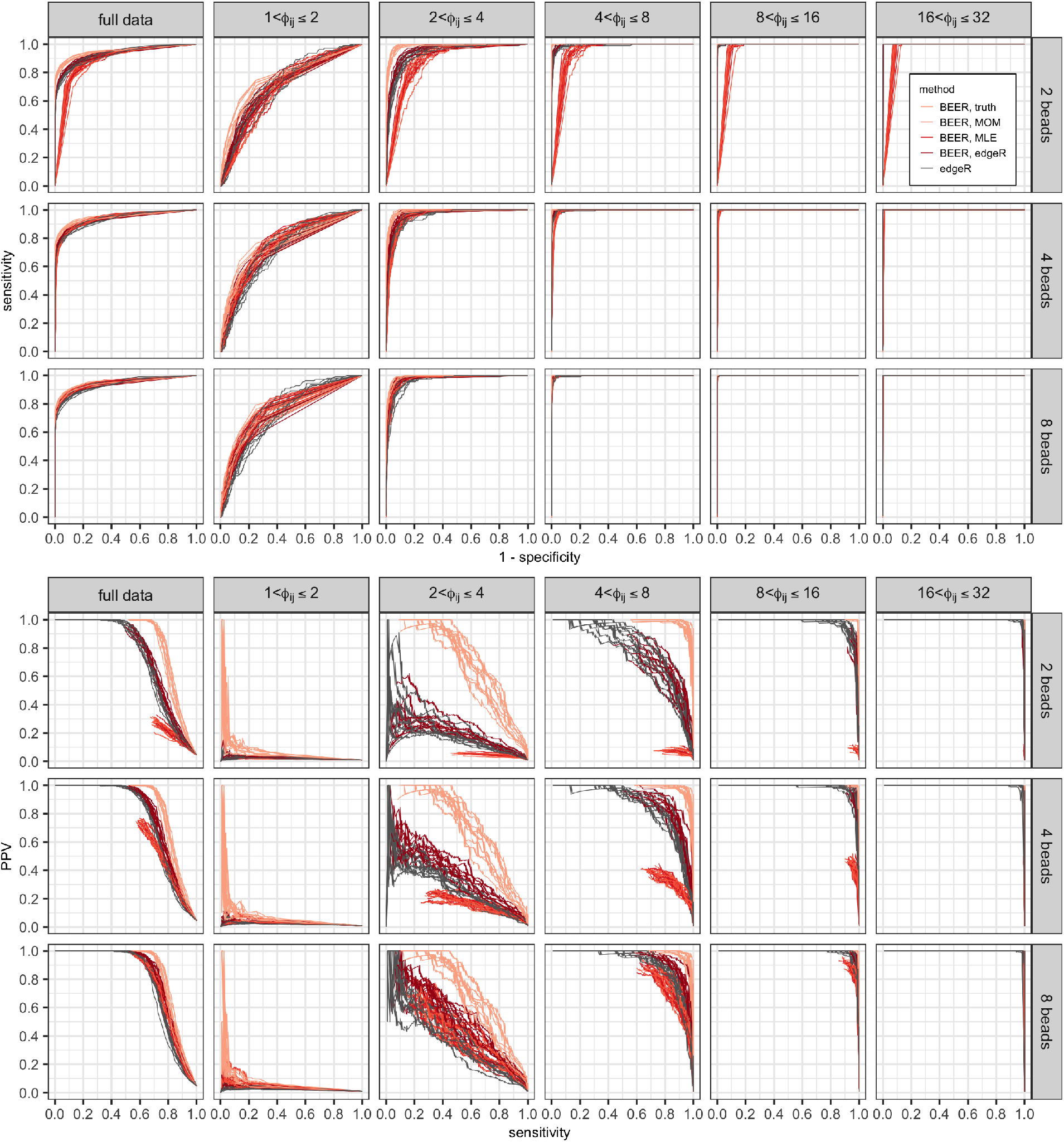
ROC (top panels) and PR (bottom panels) curves for various fold-change bins, by approach and method of estimation for *a*_*i*0_ and *b*_*i*0_.

**Figure S2:**
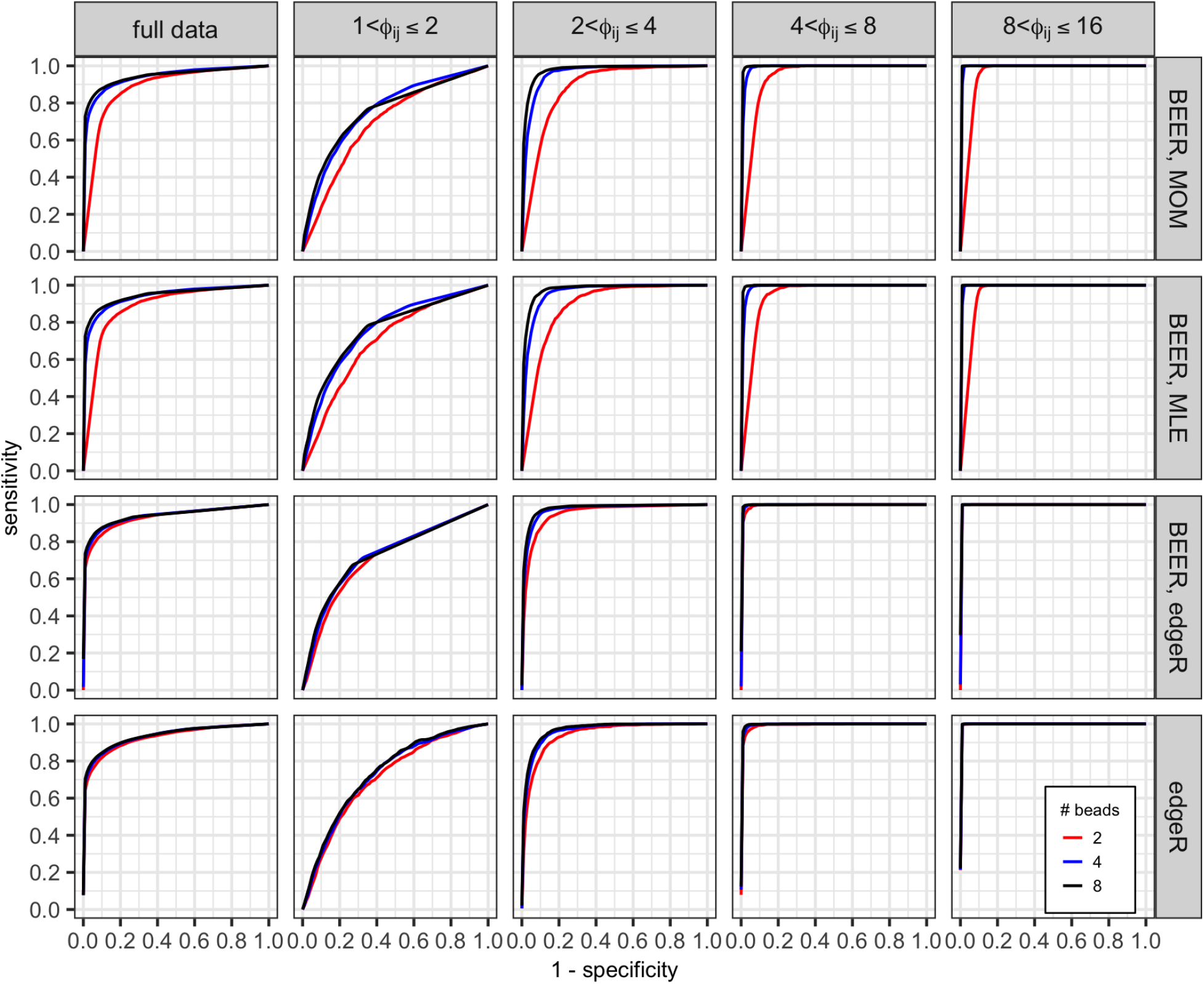
Averaged ROC curves comparing the performance of each method using 2 (red), 4 (blue), and all 8 (black) beads-only samples.

**Figure S3:**
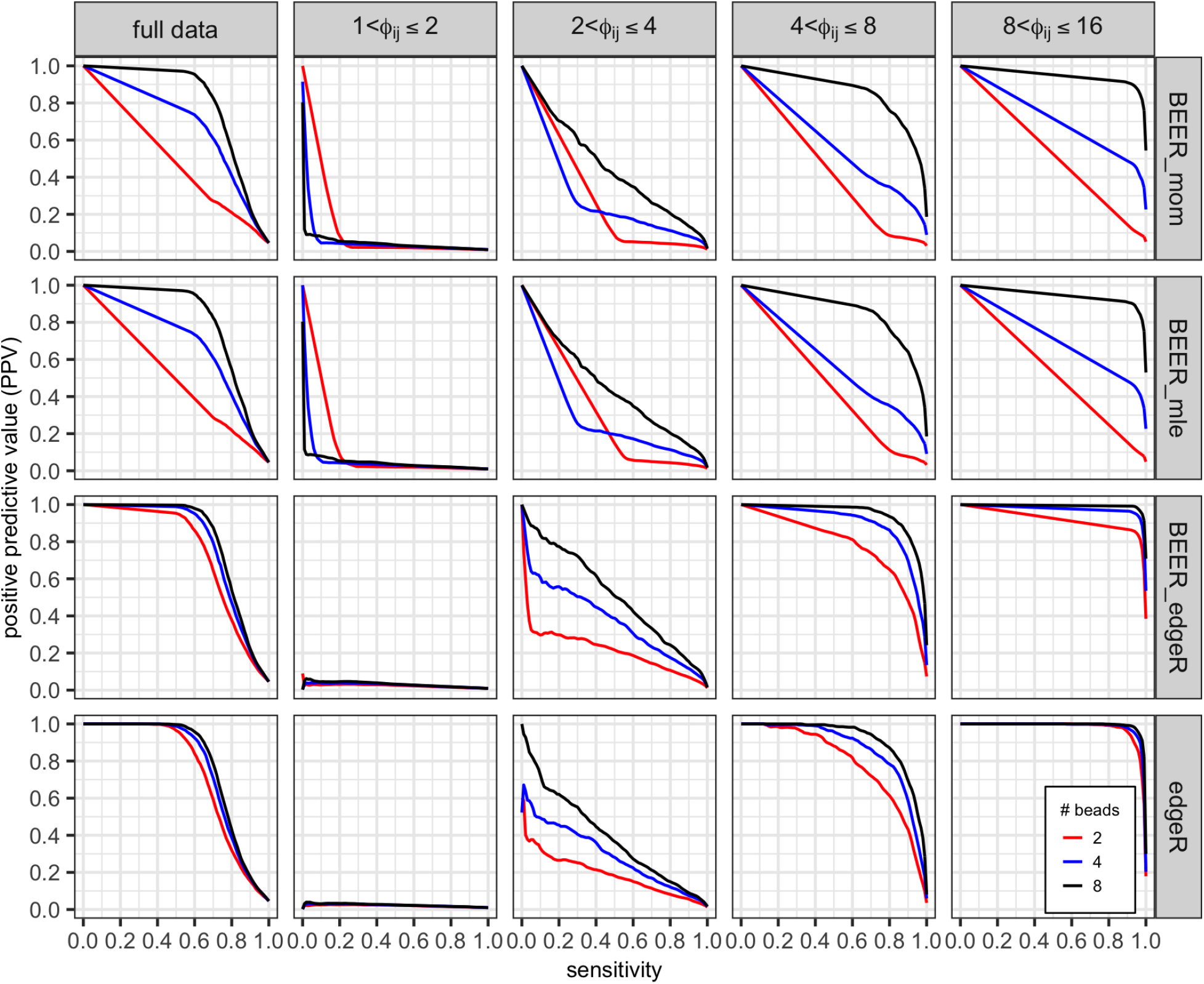
Averaged PR curves comparing the performance of each method using 2 (red), 4 (blue), and all 8 (black) beads-only samples.

**Figure S4:**
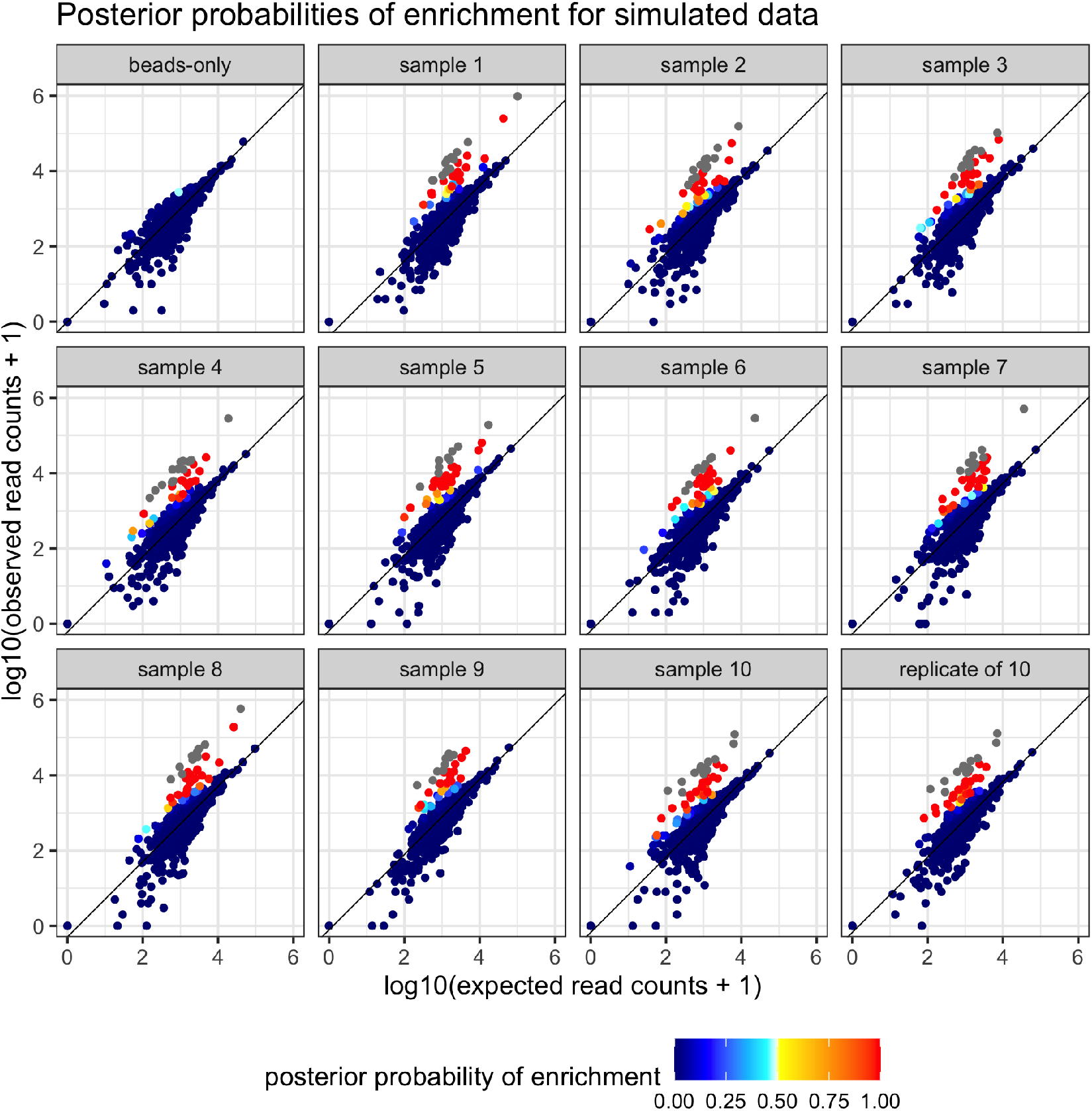
Posterior probability of enrichment for one simulated data set. Expected read counts for each peptide are derived by taking the average proportion of reads pulled in beads-only samples and multiplying the proportion by the library size of the sample. Peptides categorized as highly enriched are colored in grey. Warmer colors indicate that the peptide has over a 50% chance of being enriched. Points are plotted such that points with posterior probabilities closer to 0.5 are on top. The beads-only sample in the top left is run as a serum sample.

**Figure S5:**
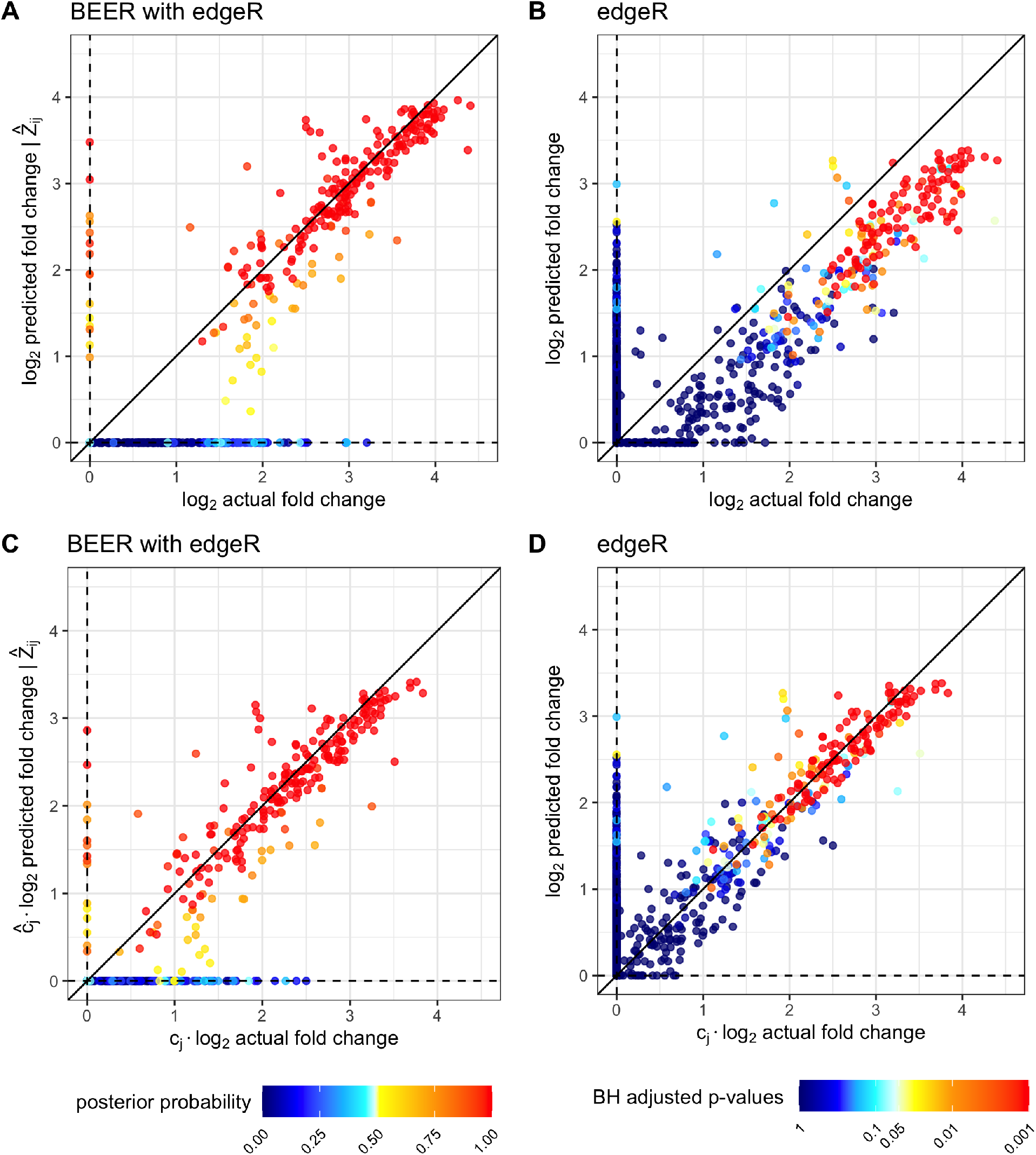
Comparison of estimated fold-changes to true fold-changes (A-B) and estimated fold-changes versus true fold-changes after adjusting for the attenuation constant (C-D) for one simulated data set. Only peptides from serum samples are included in each plot, and each peptide is represented by a point. Note that by construction, there are 120 peptides between each log_2_ increment, and highly enriched peptides are not included in the above plots. Warm colors indicate high probability of enrichment (posterior probability of enrichment > 0.5 or – log_10_(p-value) > 20).

**Figure S6:**
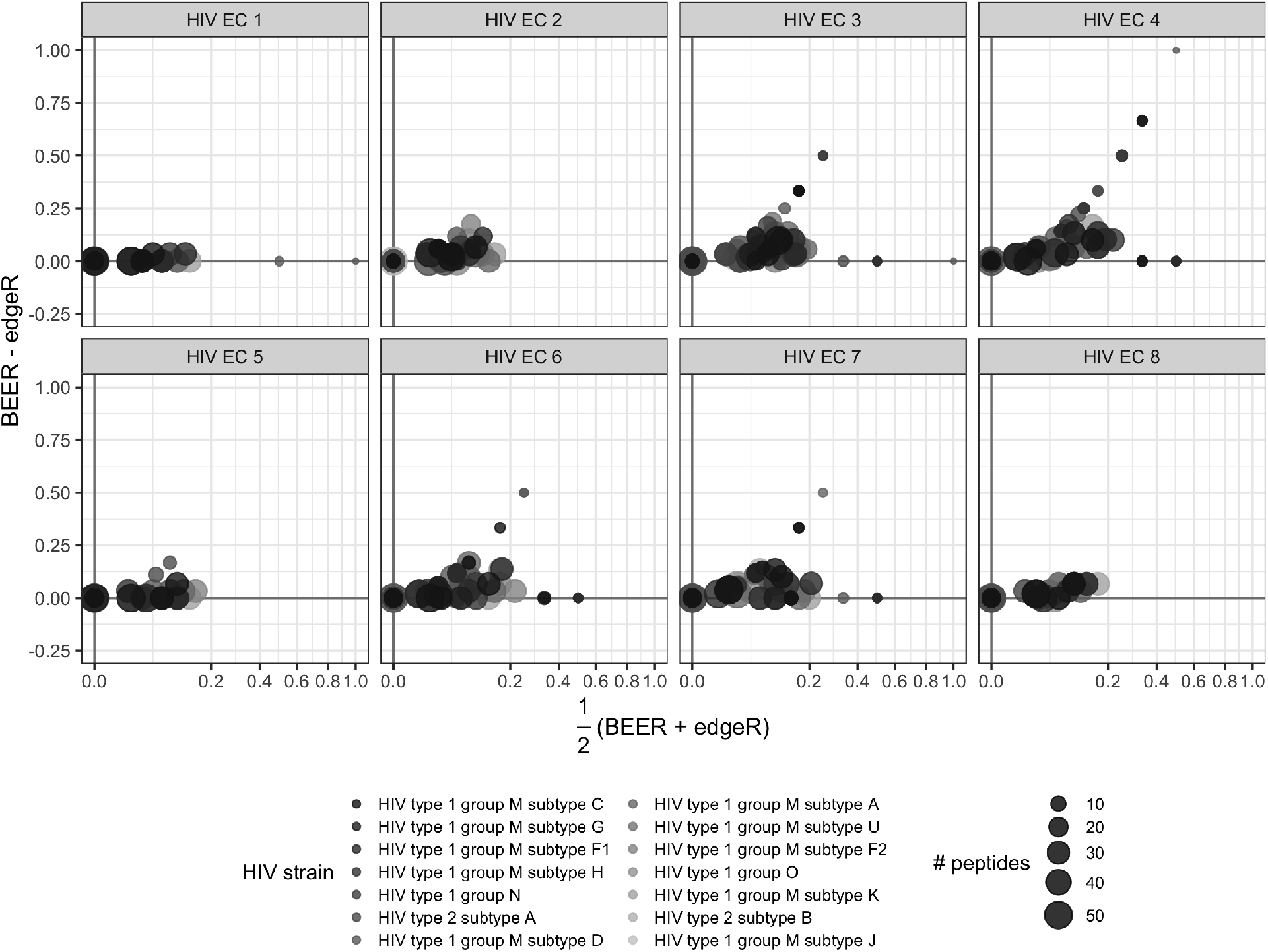
Proportion of enriched peptides by protein without HIV subtype B. Each point represents a protein. The color of the point indicates which virus the protein belongs to, and the size of the point corresponds to the number of peptides tiling the protein.

**Figure S7:**
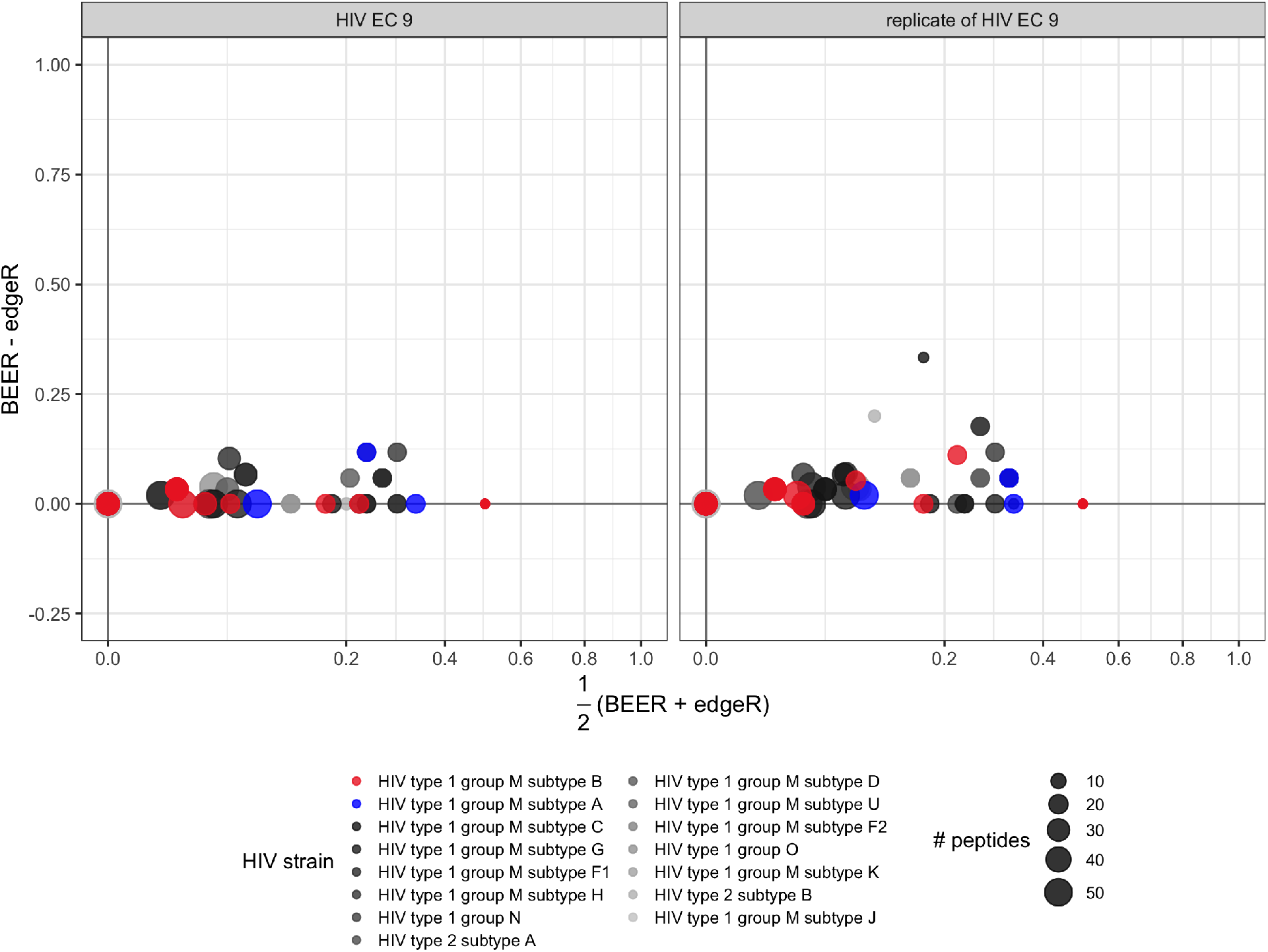
Proportion of enriched peptides by protein across technical replicates. This individual was infected with HIV subtype A. Each point represents a protein. The color of the point indicates which virus the protein belongs to, and the size of the point corresponds to the number of peptides tiling the protein.

**Figure S8:**
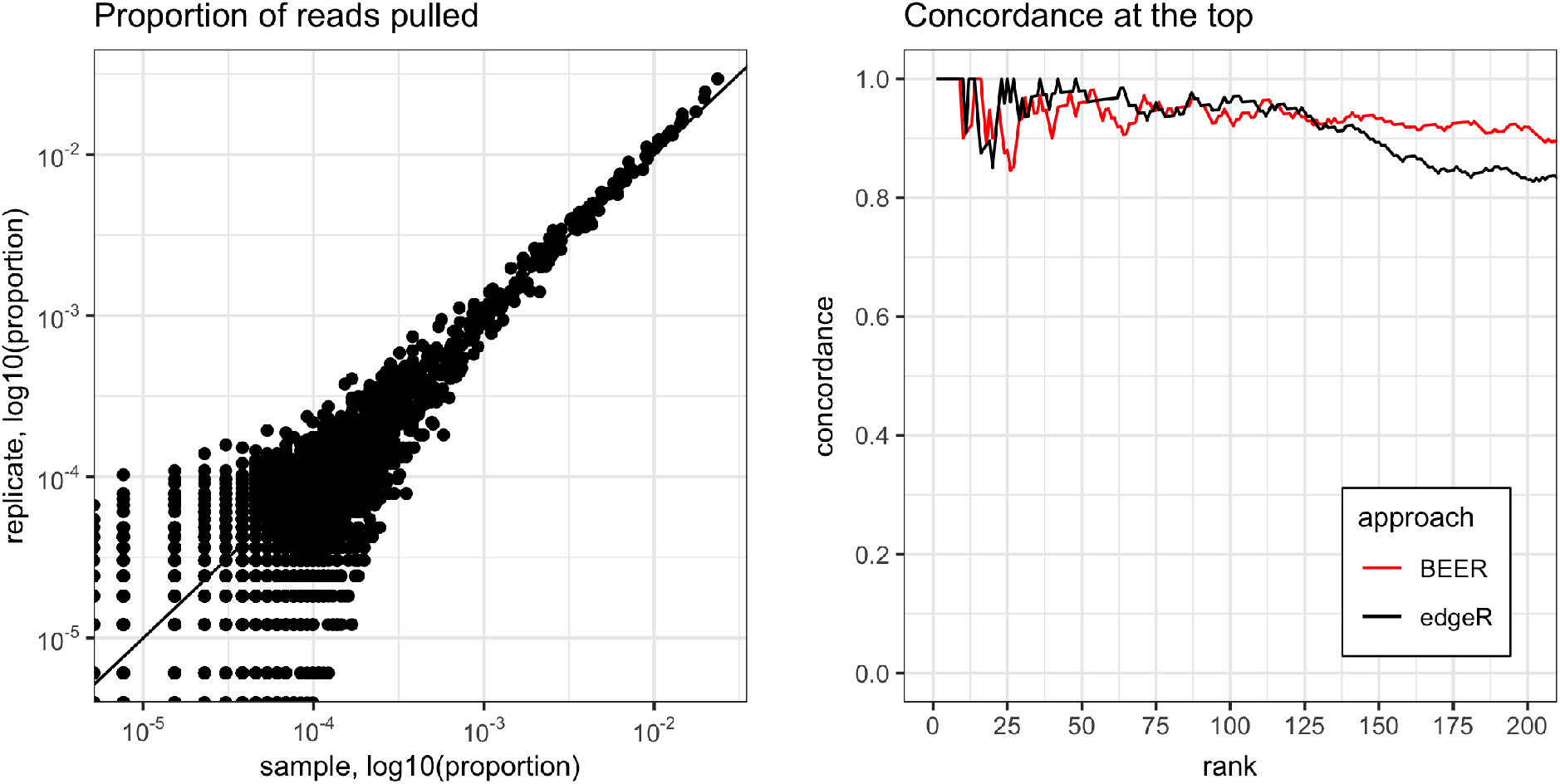
Left: proportion of reads pulled for 3,395 HIV peptides for two technical replicates. Right: concordance of HIV technical replicates, shown as proportion of peptides among the top ranked peptides in both replicates. For BEER, peptides are ranked by decreasing posterior probability of enrichment. For edgeR, peptides are ranked by increasing p-values. For both methods, ties of posterior probabilities and p-value (e.g., 0 and 1) were broken by the estimated fold-change. The top eight peptides from BEER are all highly enriched and treated exchangeably as no fold-change estimates are returned.

**Figure S9:**
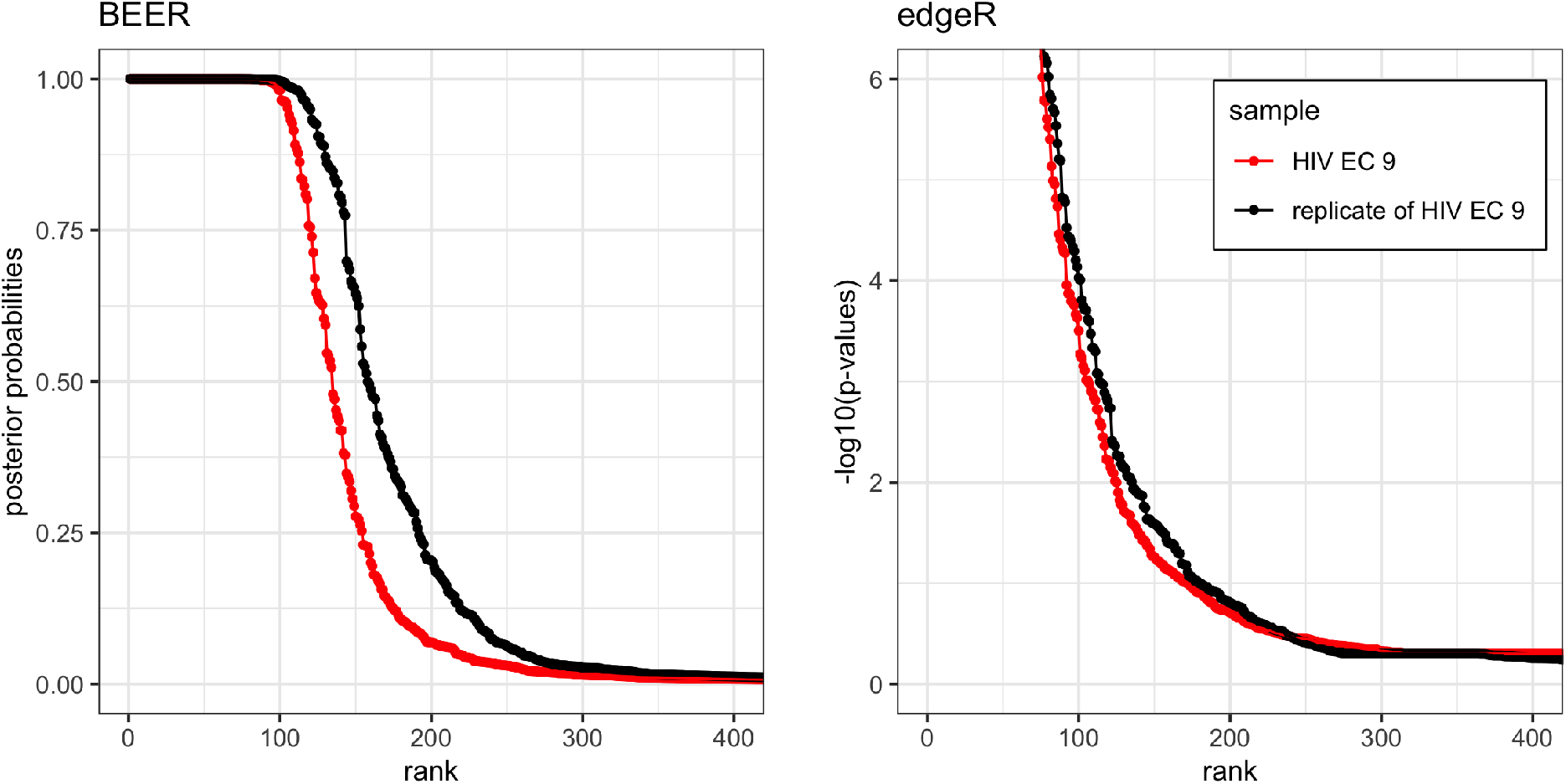
HIV replicate posterior probabilities by rank. For each of the technical replicates, peptides are sorted in decreasing order by posterior probability and −log10(edgeR p-values). For clarity of display, p-values were truncated at 10^−6^.

**Figure S10:**
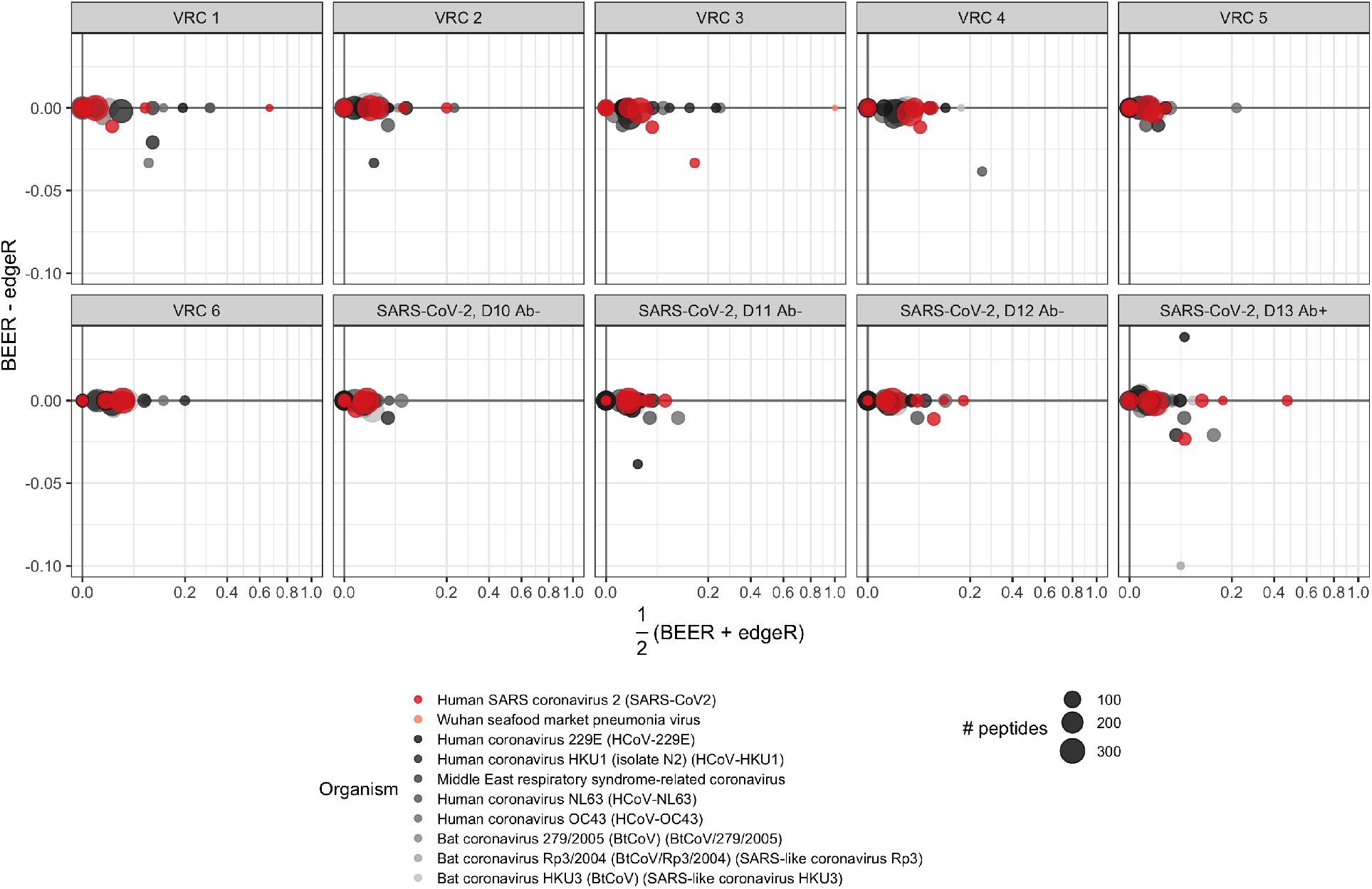
MA plots for the proportion of enriched peptides by protein for six pre-pandemic samples and four samples from one individual infected with SARS-CoV-2. Samples from this individual were collected at various days since symptom onset (labels D10-D13) and were additionally tested for SARS-CoV-2 antibodies. Antibody test results (positive or negative) are indicated by Ab+ or Ab-, respectively. Points represent individual proteins; point colors indicate virus types; and point diameters indicate the number of peptides tiling the respective proteins. In the CoronaScan library, peptides are present in duplicate, so the number of peptides is double the number of unique peptides.

**Figure S11:**
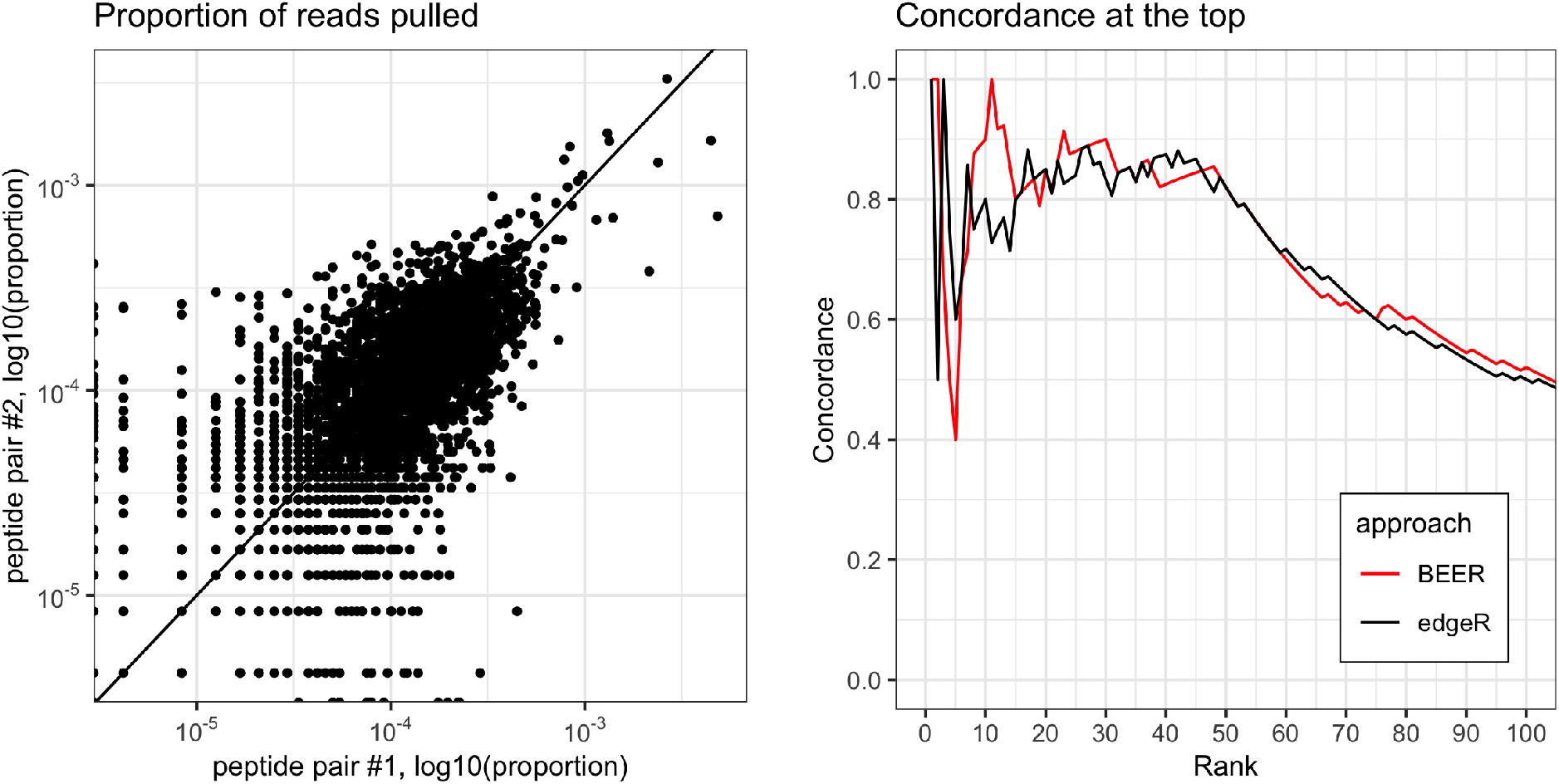
Left: concordance of paired peptides in CS sample VRC 1. For each unique peptide sequence, the proportion of reads pulled for peptide 1 with the same sequence is plotted against the proportion of reads pulled for peptide 2 of the same sequence. Right: concordance between the rankings for the top *k* ranks (x-axis) between all peptide pairs. For BEER, peptides are ranked by decreasing posterior probability of enrichment, with ties broken by the estimated fold-change (red line). For edgeR, peptides are ranked by increasing p-values, with ties again broken by estimated fold-changes (black line).

**Figure S12:**
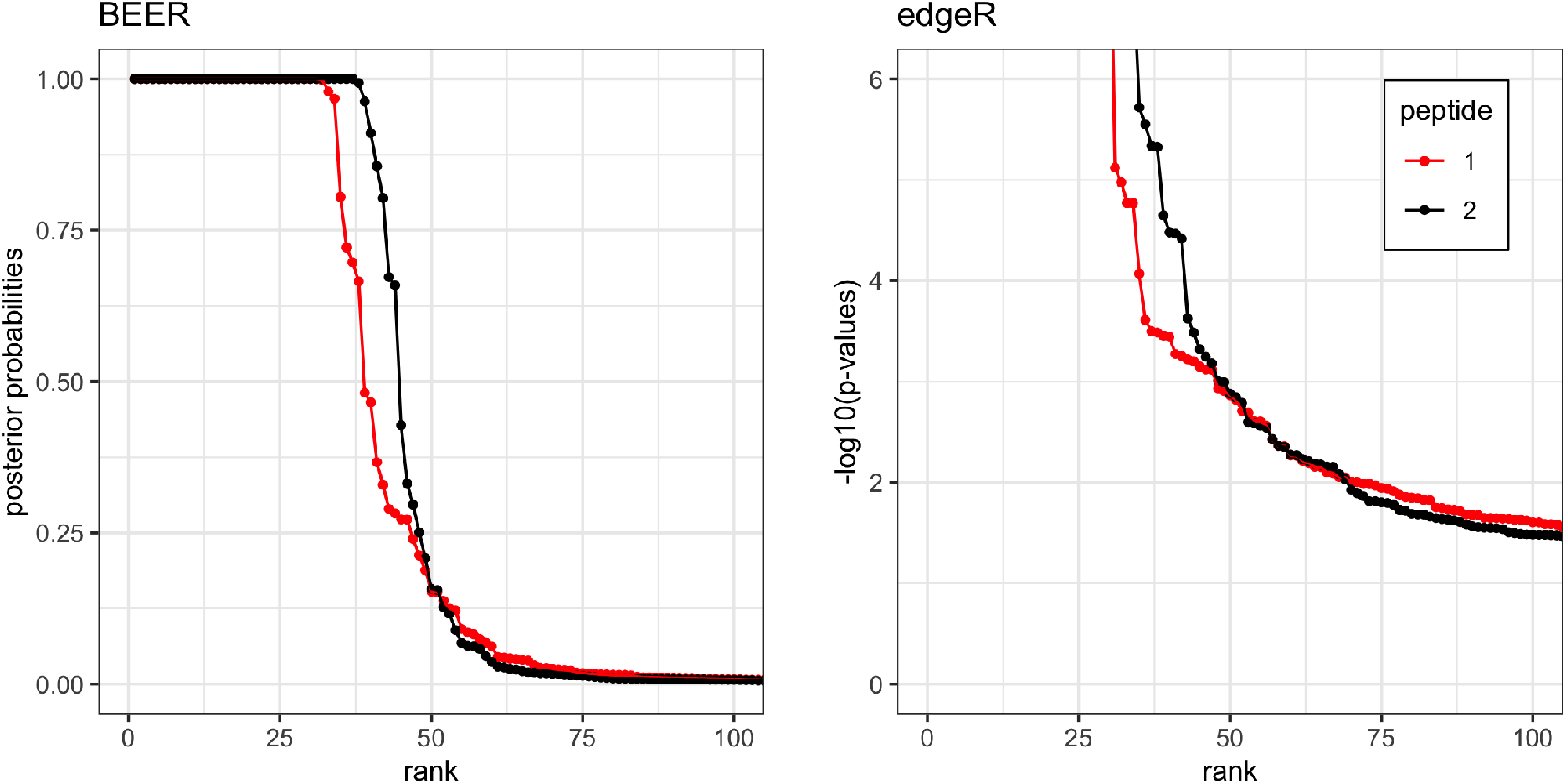
CoronaScan peptide pairs by rank. For each set of unique peptides, peptides are sorted in decreasing order by posterior probability and −log10(edgeR p-values). For clarity of display, p-values were truncated at 10^−6^.

**Figure S13:**
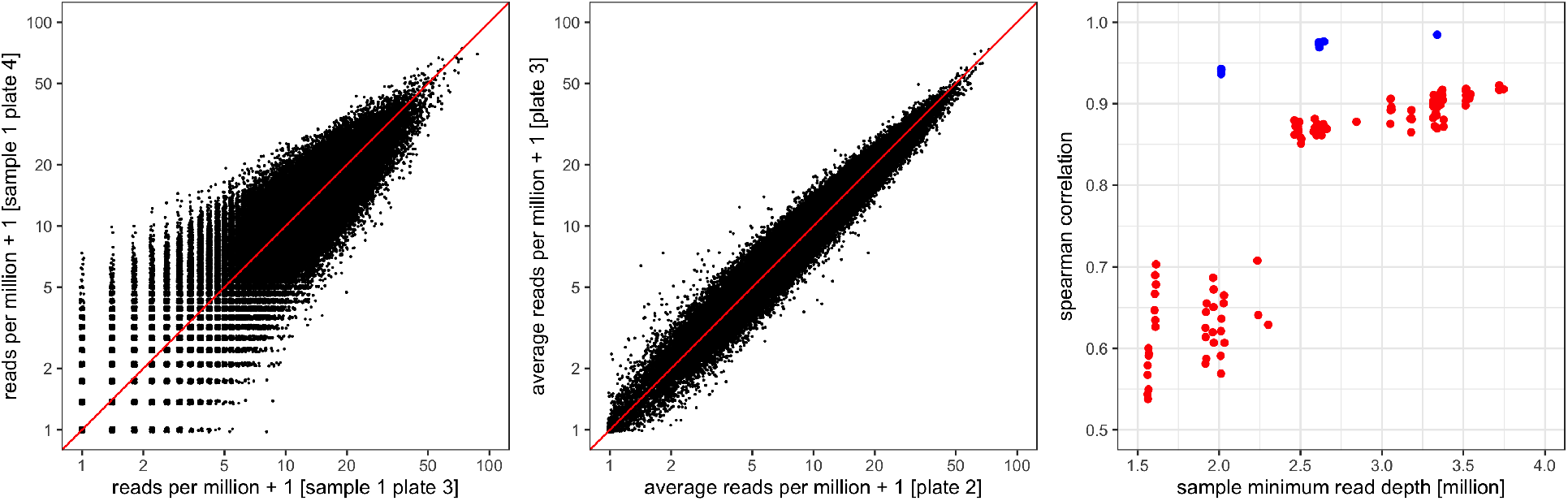
Evidence for a strong peptide effect in PhIP-Seq data, demonstrated using data from five plates of a previous experiment using HIV samples, analyzed in Eshleman et al.^18^ and Chen et al.^33^ Left: observed read counts per million reads for 95,242 peptides from two “beads only” samples from different plates. For these two samples, the Spearman correlation is 0.875. Middle: observed average read counts per million reads for 95,242 peptides from all “beads only” samples from two different plates. For these two plates, the Spearman correlation is 0.975. Right: within and between plate sample correlations as a function of sequencing depth. For each pair of bead only samples from the same plate (red dots, 117 pairs total), the Spearman correlation (y-axis) is related to the minimum of the respective two sequencing depths (x-axis). For each pair of plates (blue dots, 10 pairs total), the Spearman correlation between the average read counts of the bead only samples (y-axis) is also related to the minimum of the two median sequencing depths (x-axis), and substantially higher than the correlations of the bead only samples.

**Figure S14:**
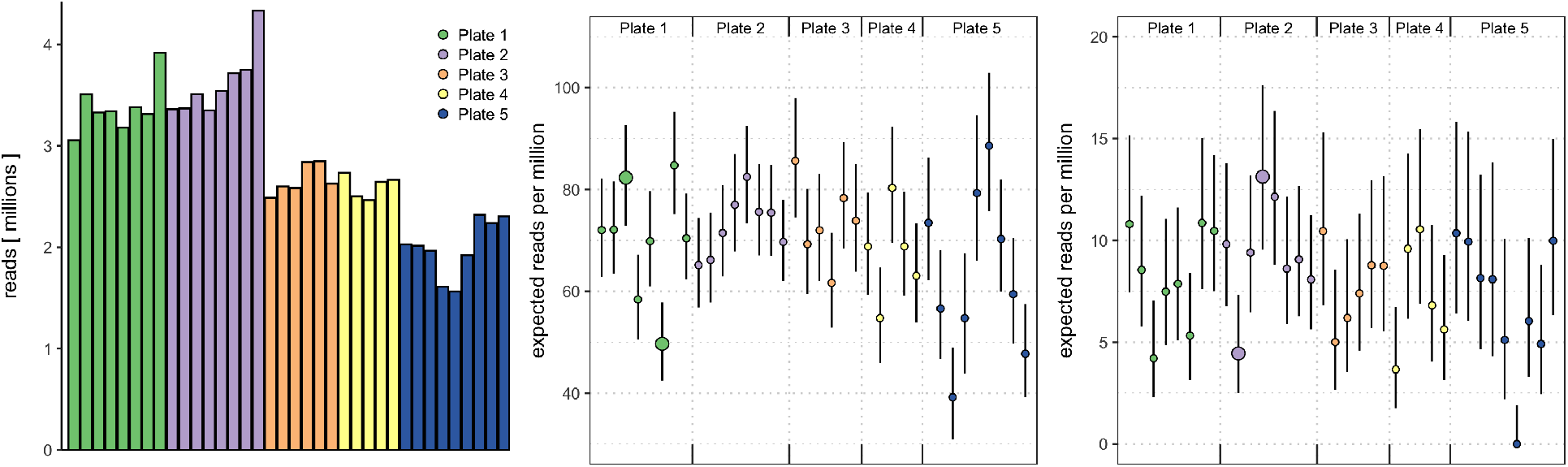
Evidence for larger than binomial variability in the PhIP-Seq data analyzed in Eshleman et al.^18^ and Chen et al.^33^ Left: library size (reads in millions) for 36 control (“bead only”) samples from 5 plates. Middle: expected read counts per million reads aligned based on the estimated Binomial probabilities (dots, colored by plate) and respective 95% confidence intervals for a peptide with large expected read counts, for each of the control samples. Highlighted are samples 3 and 6 from plate 1, showing large discrepancies between the Binomial probabilities for this peptide between the two bead only samples. Right: the same statistics as in the middle panel, for a peptide with smaller expected read counts. Highlighted are samples 10 and 12 from plate 2, again showing a large discrepancy between the Binomial probabilities for this peptide between the two bead only samples.

**Figure S15:**
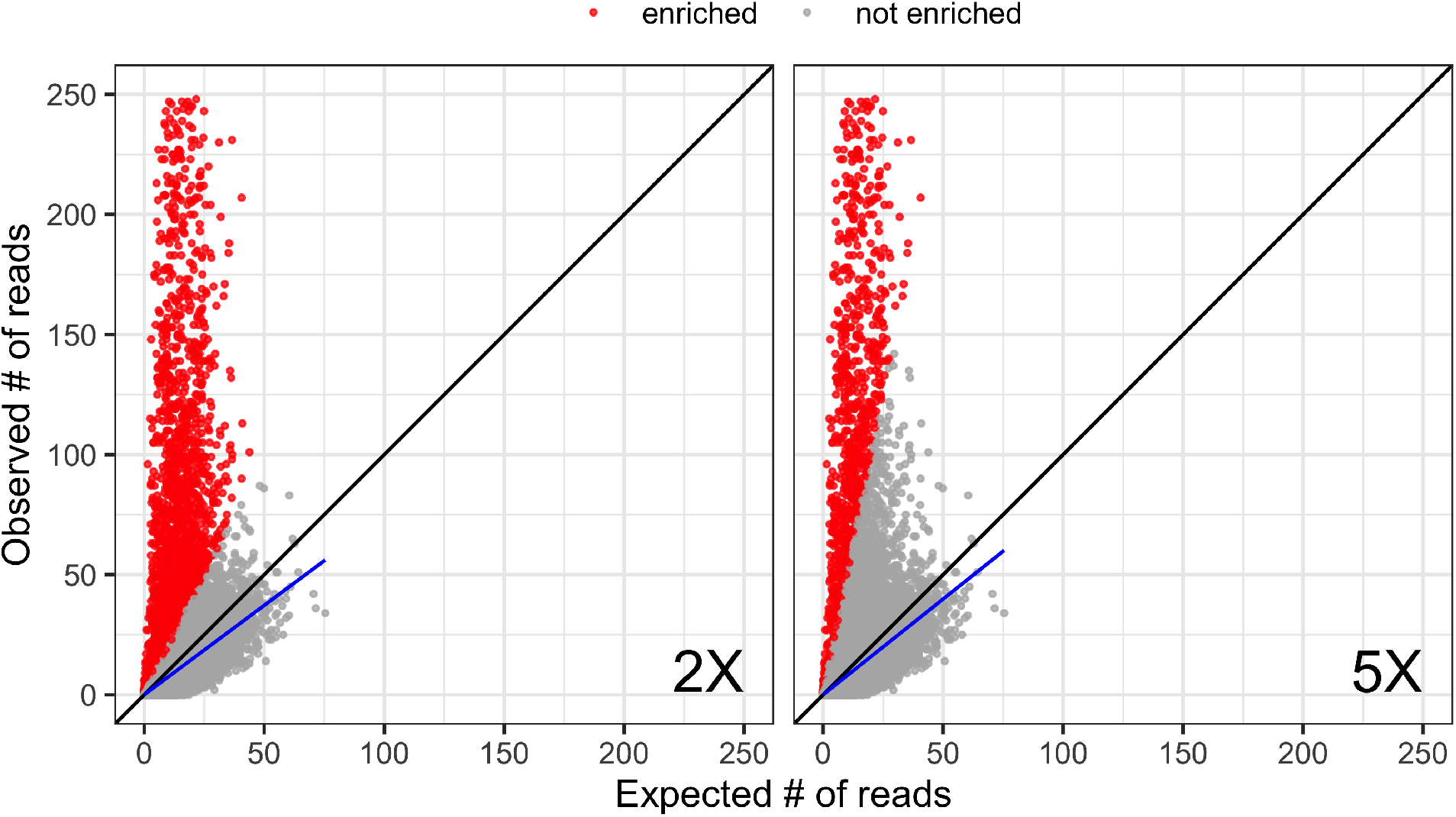
Expected versus observed read counts for 95,242 peptides from a randomly selected serum sample. Expected read counts for each peptide were derived using maximum likelihood estimates from the negative controls on the same plate. Each point represents one peptide from one sample, and peptides were considered enriched (red) if the observed read count was over 2 times (left) and 5 times (right) the expected number of reads. Linear regression lines (blue) were fitted using the non-enriched peptides and compared to the line where observed and expected reads are equal (black). The observed read counts were truncated at 250 to enhance the display.

**Figure S16:**
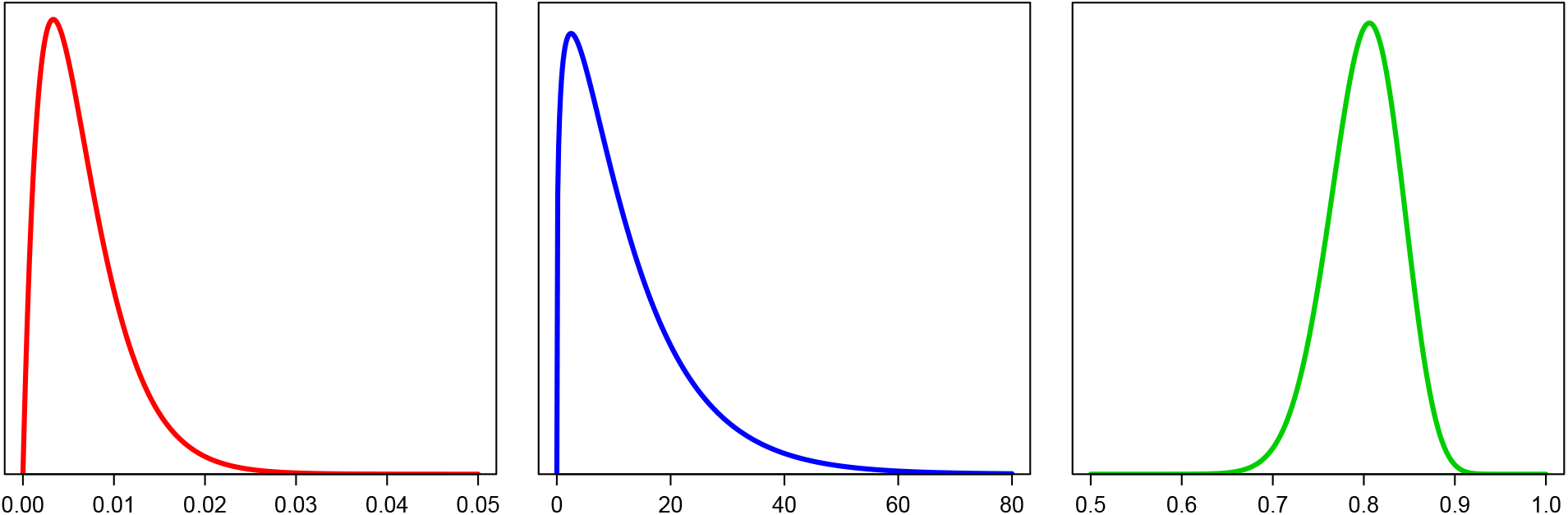
Left: the prior distribution for the proportion of reactive peptides in sample *j*, *π_j_*, modeled as a Beta distribution Beta(*a_π_* = 2, *b_π_* = 300), reflecting peptide enrichment seen in previous studies. Middle: a Gamma(*a_ϕ_* = 1.25, *b_ϕ_* = 0.1) distribution, used in the prior distribution for the fold change *ϕ_ij_* for peptide *i* in sample *j*, if reactive. Right: the prior distribution for the scaling constant in sample *j*, *c_j_*, modeled as a Beta distribution Beta(*a_c_* = 80, *b_c_* = 20).

## References

[1] Robinson MD, McCarthy DJ, Smyth GK. edgeR: a Bioconductor package for differential expression analysis of digital gene expression data. Bioinformatics, 26(1):139–140 (2010).

[2] Mohan D, Wansley DL, Sie BM, Noon MS, Baer AN, et al. PhIP-Seq characterization of serum antibodies using oligonucleotide-encoded peptidomes. Nature Protocols, 13(9):1958–1978 (2018).

[3] Larman HB, Zhao Z, Laserson U, Li MZ, Ciccia A, et al. Autoantigen discovery with a synthetic human peptidome. Nature Biotechnology, 29(6):535–541 (2011).

[4] Xu GJ, Kula T, Xu Q, Li MZ, Vernon SD, et al. Comprehensive serological profiling of human populations using a synthetic human virome. Science, 348(6239) (2015).

[5] Monaco DR, Sie BM, Nirschl TR, Knight AC, Sampson HA, et al. Profiling serum antibodies with a pan allergen phage library identifies key wheat allergy epitopes. Nature communications, 12:379 (2021).

[6] Angkeow JW, Monaco DR, Chen A, Venkataraman T, Jayaraman S, et al. Prevalence, persistence, and genetics of antibody responses to protein toxins and virulence factors. Biorxiv https://www.biorxivorg/content/101101/20211001462481v1 (2021).

[7] Morgenlander WR, Henson SN, Monaco DR, Chen A, Littlefield K, et al. Antibody responses to endemic coronaviruses modulate COVID-19 convalescent plasma functionality. The Journal of clinical investigation, 131 (2021).

[8] Larman HB, Laserson U, Querol L, Verhaeghen K, Solimini NL, et al. PhIP-Seq characterization of autoantibodies from patients with multiple sclerosis, type 1 diabetes and rheumatoid arthritis. Journal of autoimmunity, 43:1–9 (2013).

[9] Larman HB, Salajegheh M, Nazareno R, Lam T, Sauld J, et al. Cytosolic 5’-nucleotidase 1A autoimmunity in sporadic inclusion body myositis. Annals of neurology, 73:408–418 (2013).

[10] Xu GJ, Shah AA, Li MZ, Xu Q, Rosen A, et al. Systematic autoantigen analysis identifies a distinct subtype of scleroderma with coincident cancer. Proceedings of the National Academy of Sciences of the United States of America, 113:E7526–E7534 (2016).

[11] Pou C, Nkulikiyimfura D, Henckel E, Olin A, Lakshmikanth T, et al. The repertoire of maternal anti-viral antibodies in human newborns. Nature medicine, 25:591–596 (2019).

[12] Isnard P, Kula T, Avettand Fenoel V, Anglicheau D, Terzi F, et al. Temporal virus serological profiling of kidney graft recipients using VirScan. Proceedings of the National Academy of Sciences of the United States of America, 116:10899–10904 (2019).

[13] Finton KAK, Larimore K, Larman HB, Friend D, Correnti C, et al. Autoreactivity and exceptional CDR plasticity (but not unusual polyspecificity) hinder elicitation of the anti-HIV antibody 4E10. PLoS pathogens, 9:e1003639 (2013).

[14] Finton KAK, Friend D, Jaffe J, Gewe M, Holmes MA, et al. Ontogeny of recognition specificity and functionality for the broadly neutralizing anti-HIV antibody 4E10. PLoS pathogens, 10:e1004403 (2014).

[15] Kammers K, Chen A, Monaco DR, Hudelson SE, Grant-McAuley W, et al. HIV Antibody Profiles in HIV Controllers and Persons With Treatment-Induced Viral Suppression. Frontiers in immunology, 12:740395 (2021).

[16] Schubert RD, Hawes IA, Ramachandran PS, Ramesh A, Crawford ED, et al. Pan-viral serology implicates enteroviruses in acute flaccid myelitis. Nature medicine, 25:1748–1752 (2019).

[17] Mina MJ, Kula T, Leng Y, Li M, de Vries RD, et al. Measles virus infection diminishes preexisting antibodies that offer protection from other pathogens. Science (New York, NY), 366:599–606 (2019).

[18] Eshleman SH, Laeyendecker O, Kammers K, Chen A, Sivay MV, et al. Comprehensive Profiling of HIV Antibody Evolution. Cell reports, 27:1422–1433.e4 (2019).

[19] Vogl T, Klompus S, Leviatan S, Kalka IN, Weinberger A, et al. Population-wide diversity and stability of serum antibody epitope repertoires against human microbiota. Nature medicine, 27:1442–1450 (2021).

[20] Venkataraman T, Valencia C, Mangino M, Morgenlander W, Clipman SJ, et al. Analysis of antibody binding specificities in twin and SNP-genotyped cohorts reveals that antiviral antibody epitope selection is a heritable trait. Immunity, 55(1):174–184.e5 (2022).

[21] McCarthy DJ, Chen Y, Smyth GK. Article Navigation Differential expression analysis of multifactor RNA-Seq experiments with respect to biological variation. Nucleic Acids Research, 40(10):4288–4297 (2012).

[22] Love MI, Huber W, Anders S. Moderated estimation of fold change and dispersion for RNA-seq data with DESeq2. Genome Biology, 15(12):550 (2014).

[23] Law CW, Chen Y, Shi W, Smyth GK. voom: precision weights unlock linear model analysis tools for RNA-seq read counts. Genome Biology, 15(2):R29 (2014).

[24] Robinson MD, Oshlack A. A scaling normalization method for differential expression analysis of RNA-seq data. Genome biology, 11:R25 (2010).

[25] Hansen KD, Wu Z, Irizarry RA, Leek JT. Sequencing technology does not eliminate biological variability. Nature biotechnology, 29:572–573 (2011).

[26] Nocedal J, Wright S. Numerical Optimization. Springer New York (1999).

[27] Anders S, Huber W. Differential expression analysis for sequence count data. Genome biology, 11:R106 (2010).

[28] Kammers K, Cole RN, Tiengwe C, Ruczinski I. Detecting Significant Changes in Protein Abundance. EuPA open proteomics, 7:11–19 (2015).

[29] Ritchie ME, Phipson B, Wu D, Hu Y, Law CW, et al. limma powers differential expression analyses for RNA-sequencing and microarray studies. Nucleic acids research, 43:e47 (2015).

[30] Plummer M. JAGS: A program for analysis of Bayesian graphical models using Gibbs sampling (2003).

[31] R Core Team. R: A Language and Environment for Statistical Computing. R Foundation for Statistical Computing, Vienna, Austria (2021).

[32] Plummer M. rjags: Bayesian Graphical Models using MCMC (2019). R package version 4-10.

[33] Chen A, Laeyendecker O, Eshleman SH, Monaco DR, Kammers K, et al. A top scoring pairs classifier for recent HIV infections. Statistics in medicine, 40:2604–2612 (2021).

[34] Robinson MD, Smyth GK. Small-sample estimation of negative binomial dispersion, with applications to SAGE data. Biostatistics (Oxford, England), 9:321–332 (2008).

